# 3 minutes to precisely measure morphogen concentration

**DOI:** 10.1101/305516

**Authors:** Tanguy Lucas, Huy Tran, Carmina Angelica Perez Romero, Aurélien Guillou, Cécile Fradin, Mathieu Coppey, Aleksandra M. Walczak, Nathalie Dostatni

## Abstract

Morphogen gradients provide concentration-dependent positional information along polarity axes. Although the dynamics of establishment of these gradients is well described, precision and noise in the downstream activation processes remain elusive. A simple paradigm to address these questions is the Bicoid morphogen gradient that elicits a rapid step-like transcriptional response in young fruit fly embryos. Focusing on the expression of the main Bicoid target, *hunchback* (*hb*), at the onset of zygotic transcription, we used the MS2-MCP approach which combines fluorescent labeling of nascent mRNA with live imaging at high spatial and temporal resolution. Removing 36 putative Zelda binding sites unexpectedly present in the original MS2 reporter, we show that the 750 bp of the *hb* promoter are sufficient to recapitulate endogenous expression at the onset of zygotic transcription. After each mitosis, in the anterior, expression is turned on to rapidly reach a plateau with all nuclei expressing the reporter. Consistent with a Bicoid dose-dependent activation process, the time period required to reach the plateau increases with the distance to the anterior pole. Remarkably, despite the challenge imposed by frequent mitoses and high nuclei-to-nuclei variability in transcription kinetics, it only takes 3 minutes at each interphase for the MS2 reporter loci to measure subtle differences in Bicoid concentration and establish a steadily positioned and steep (Hill coefficient ~ 7) expression boundary. Modeling based on cooperativity between the 6 known Bicoid binding sites in the *hb* promoter region and assuming rate limiting concentrations of the Bicoid transcription factor at the boundary is able to capture the observed dynamics of pattern establishment but not the steepness of the boundary. This suggests that additional mechanisms are involved in the steepness of the response.

## Introduction

Morphogens are at the origin of complex axial polarities in many biological systems. In these systems, positional information is proposed to be provided by the morphogen concentrations, which allow each cell to measure its position along the axis and turn on expression of target genes responsible for the determination of its identity. Although the existence of these gradients is now well established (1), the quantitative details of their functioning (i.e. how small differences in morphogen concentrations are precisely and robustly interpreted into a threshold-dependent step-like response) remains largely debated (2).

To address this question, we study the Bicoid morphogen, which specifies cell identity along the antero-posterior (AP) axis of the fruit fly embryo (3). The Bicoid concentration gradient culminates at the anterior pole (4) and is distributed steadily in an exponential gradient along the AP axis after one hour of development (4–6) (Fig 1A). Bicoid is a homeodomain transcription factor which binds DNA. Bicoid binding sites are found in the regulatory sequences of Bicoid target genes and are both necessary (7–9) and sufficient (10–12) for Bicoid-dependent expression. Changes in Bicoid dosage induce a shift of their expression domain along the AP axis (9, 13) indicating that Bicoid provides concentration-dependent positional information to the system (14). Remarkably, just 30 min after the onset of zygotic transcription and the steady establishment of the Bicoid gradient, the main Bicoid target gene, *hunchback* (hb), already exhibits a step-like expression pattern. This pattern is characterized by an anterior domain containing almost exclusively *hb* transcriptionally active nuclei and a posterior domain containing exclusively *hb* silent nuclei (3) (Fig 1B). Remarkably, the boundary separating expressing and non-expressing nuclei is very steep despite the stochastic nature of transcription in eukaryotic cells (15), the short interphase duration (~5 min) and the subtle difference in the Bicoid concentration (10%) on either side of the boundary (16). How the *hb* pattern so rapidly acquires such a steep boundary with high levels of expression in the whole anterior domain is unclear. It could involve a purely quantitative threshold-dependent process in which Bicoid, acting as a direct transcription activator, is the main source of positional information. Alternatively, additional mechanisms including posterior inhibitors such as those acting later during development at cycle 14 (7, 17) might be at work. In any case, the mechanism involved should account for a Bicoid dose-dependent effect on the positioning of the boundary along the AP axis. Also, given the early timing of development, most actors in this process are likely to be already present at the onset of zygotic transcription and therefore maternally provided.

**Fig 1:**
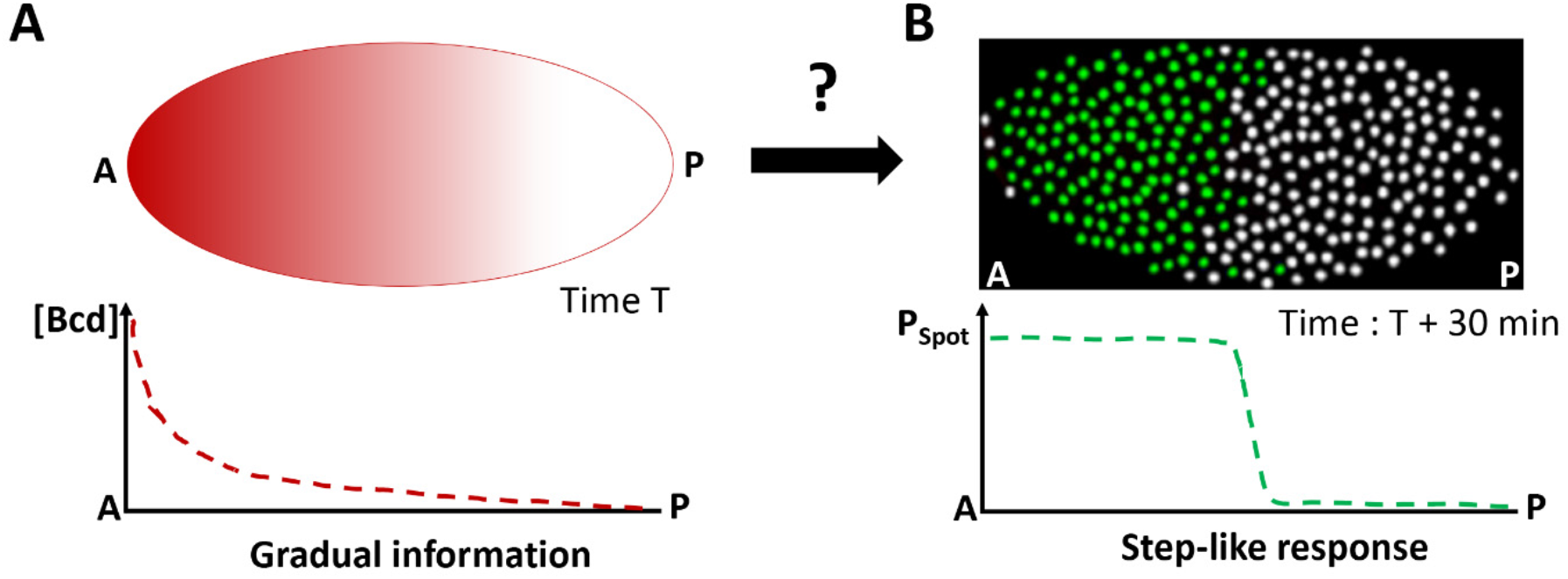
The Bicoid system transforms the gradual information contained in the Bicoid concentration gradient into a step-like response in 30 min. **A)** At nc 8 (T = 1 hr), the Bicoid exponential gradient (red) is steadily established (5, 6) with its highest concentration at the anterior pole (A) and its lowest concentration at the posterior pole (P). The first hints of zygotic transcription are detected by RNA FISH marking the onset of zygotic transcription (16). **B)** At nc 11 (T = 1 hr 30 min), the main Bicoid target gene, *hunchback (hb)*, is expressed within a large anterior expression domain. *hb* expression is schematized here from RNA FISH data (16): nuclei where ongoing transcription at the *hb* loci is detected are shown in green and nuclei silent for *hb* are shown in white.

To shed light on the formation of the *hb* expression boundary during these early steps of development, we have previously adapted the MS2-MCP approach to developing fly embryos (18). This approach allows the fluorescent tagging of RNA in living cells and provides access to the transcription dynamics of a MS2 reporter locus (19, 20). In our first attempt, we placed the *hb* canonical promoter region (~ 750 bp) upstream of an MS2 cassette containing 24 MS2 loops and analyzed expression of this reporter *(hb-MS2)* in embryos expressing the MCP-GFP protein maternally (18). The *hb-MS2* reporter was expressed very early as robustly as endogenous *hb* in the anterior half of the embryo. However, unlike endogenous *hb*, the reporter was also expressed in the posterior, though more heterogeneously and more transiently than in the anterior. The different expression of the *hb-MS2* reporter and endogenous *hb* in the posterior suggested that, the *hb* canonical promoter was not sufficient to recapitulate the endogenous expression of *hb* at the onset of zygotic transcription. The most obvious interpretation of these discrepancies was that the *hb-MS2* reporter was missing key cis-response elements allowing repression of endogenous *hb* by an unknown repressor in the posterior (18).

Here, we first show that the homogenously distributed Zelda transcription factor is responsible for the expression of the *hb-MS2* reporter in the posterior. BAC recombineering indicated that the MS2 cassette itself mediates Zelda posterior expression and *in silico* analysis reveals the presence of about 36 putative Zelda binding sites in the MS2 cassette. A new reporter *(hb-MS2ΔZelda)*, placing the canonical *hb* promoter upstream of a new MS2 cassette (MS2-ΔZelda) in which those unfortunate Zelda binding sites have been mutated, faithfully recapitulates the early expression of endogenous *hb* observed by RNA FISH. Thus, the *hb* canonical promoter is sufficient for a robust step-like expression of the reporter. Quantitative analysis of the MS2 time traces of this new reporter reveals a transcription process dividing the anterior of the embryo in a saturating zone with stable features and a limiting zone closer to the boundary, with more variable features. A high probability for the promoter to be ON (Pon) is reached faster in the anterior where the concentration of Bicoid is higher than close to the boundary where Bicoid concentration is lower. Remarkably, in each interphase, full step-like response is established in not more than three minutes after which the expression boundary is locked at a given position along the AP axis. To understand this observed dynamics, we used a simple model of position readout through the binding/unbinding of a transcription factor to *N* operator sites on the *hb* promoter (21). The model that best fits the data is able to capture the very fast dynamics of establishment of the boundary. However, high steepness of the experimental pattern (coefficient of the fitted Hill function *N_Hill_* ~ 7, based on various features of the MS2 time-traces) is not achievable assuming the presence of only *N*=6 Bicoid binding sites, which is the number of known Bicoid binding sites in the canonical *hb* promoter (8). This indicates the requirement for additional mechanisms to define the steepness of the boundary.

## Results

### Zelda induces posterior expression of the hb-MS2 transgene at early cycles

The Zelda transcription factor is the major regulator of the first wave of zygotic transcription in fruit fly embryos (22) and is involved in the transcriptional regulation of the *hb* gene (23). To determine how Zelda contributes to the expression of the *hb-MS2* reporter, we analyzed expression of the reporter by live imaging (SI, Movie1) and double RNA FISH (using *hunchback* and *MS2* probes, Fig 2A) in embryos expressing various amounts of maternal Zelda. As expected (18), wild-type embryos show high expression of the *hb-MS2* reporter both in the anterior and the posterior domain ((18) and Fig 2B, top panel), with an average of 80% of expressing nuclei dispersed through the posterior domain (Fig 2B, bottom panel). In embryos from *zelda* heterozygous mutant females (*zld^mat-/+^*), expression of the *hb-MS2* reporter is reduced by 5 fold in the posterior (with only 15% of active nuclei). In embryos from *zelda* mutant germline clones, completely devoid of Zelda maternal contribution (*zld^mat-/+^*), expression boundaries of endogenous *hb* and *hb-MS2* reporter are shifted towards the anterior (Fig 2D). Moreover, posterior expression of the *hb-MS2* reporter is reduced to less than 1% of posterior nuclei (Fig 2D). We confirm thus that Zelda maternal proteins contribute to the early expression the endogenous *hb* (23) and conclude that posterior expression of the *hb-MS2* reporter in early embryos is mostly due to Zelda.

**Fig 2:**
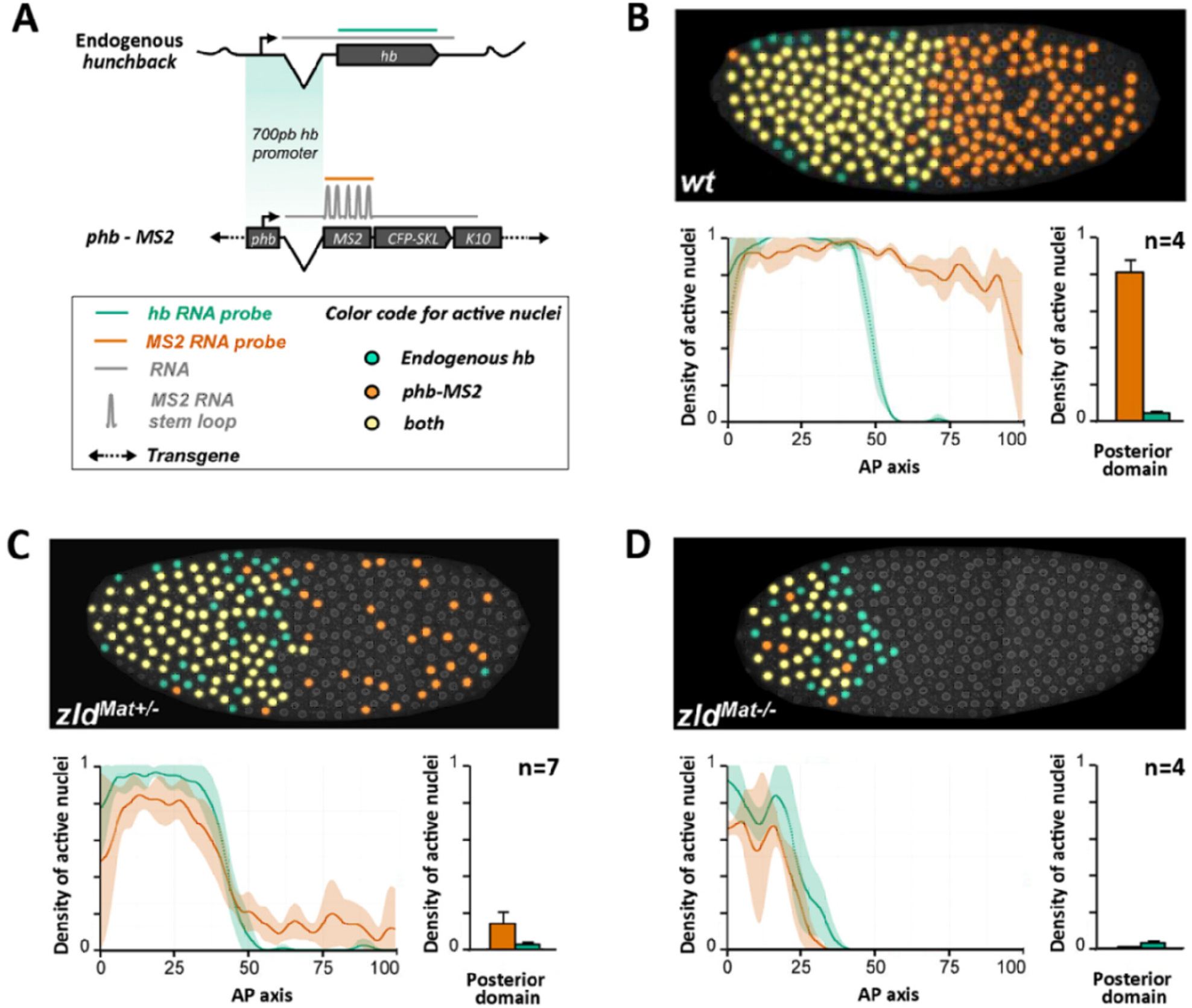
posterior expression of the *hb-MS2* reporters is Zelda dependent. **A)** Dual RNA FISH using an *hb* probe (green) and a *MS2* (orange) probe on wild-type embryos carrying one copies of the *hb-MS2* reporter, placing the *MS2-CFP-SKL-K10* cassette under the control of the 750 bp canonical promoter of *hb* (18). **B-D)** Embryos are wild-type (**B**), from heterozygous mutant females for Zelda (**C**) or germline mutant clones for Zelda (**D**). **Top panels**: Expression map of cycle 11 embryos after segmentation of nuclei and automated processing of FISH staining. Nuclei expressing only *hb* are labelled in green, nuclei expressing only the *hb-MS2* reporter are labelled in orange and nuclei expressing both *hb* and the *hb-MS2* reporter are labelled in yellow. **Bottom panels**: On the left, density of active nuclei for either *hb* (green) or the *hb-MS2* reporter (orange) along the AP axis with the anterior pole on the left (0) and the posterior pole on the right (100). On the right, density of active nuclei for the *hb-MS2* reporter (orange) and *hb* (green) in the posterior domain. For each embryo, the position of the expression boundary is calculated as the position of the maximal derivative of the active nuclei density curve. Mean were calculated for n embryos and error bars correspond to standard deviation.

### The MS2 cassette carries the cis-acting sequence responsible for hb-MS2 posterior expression

To understand the discrepancies of expression between endogenous *hb* and the *hb-MS2* reporter at early cycles, we first aimed at identifying the minimal sequence sufficient to recapitulate early expression of the *hb* locus. While a transgene carrying 18 kb of the *hb* locus was shown to recapitulate *hb* anterior expression at nc13 and nc14, its expression at earlier cycles had not been documented (24–26). RNA FISH, using a *hb* probe on embryos carrying a single insertion of the hb-18kb BAC, reveals ongoing transcription at the three hb-encoded loci and indicates that expression of the hb-18kb BAC and endogenous *hb* largely overlap in the majority of anterior nuclei (Fig 3B). Remarkably, no RNA FISH signals are detected within the posterior domain of these embryos indicating that the hb-18kb BAC encompasses all the regulatory sequences to spatially control *hb* expression during early nuclear cycles.

**Fig 3:**
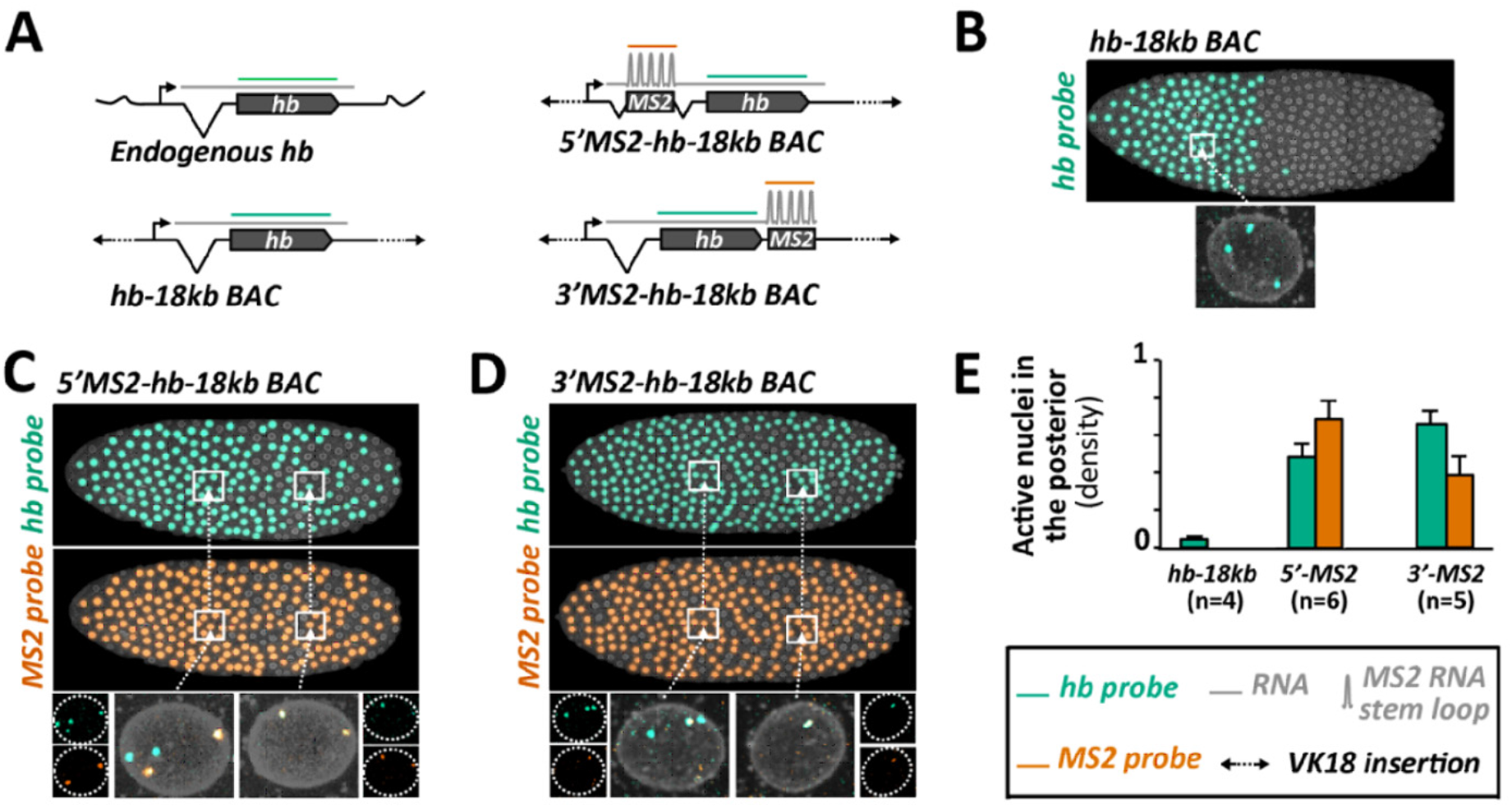
Posterior expression of the *hb-MS2* reporters is mediated *in cis* by the MS2 cassette. **A)** Structures of the *hb* expressing loci and relative positioning of the *hb* probe (green) or the *MS2* probe (orange) hybridizing to the transcripts. The *hb-18kb BAC* encompasses 18 kb of the chromosome III spanning the *hb* locus. The 1.3kb MS2 cassette is inserted in the *hb-18kb BAC* within the intron of *hb*, 0.7kb downstream the transcription start site (*5’MS2-hb-18kb* BAC, top right) or within the 3’UTR of *hb*, ~ 3 kb downstream the transcription start site (*3’MS2-hb-18kb* BAC, bottom right). All the BACs are inserted at the same position (VK18, chromosome II) within the fly genome. **B)** Top: Expression map of a typical nc11 embryo heterozygous for the *hb-18kb BAC* insertion. As highlighted on one example (bottom), nuclei in the anterior exhibit three sites of ongoing transcription detected with the *hb* probe: two of these sites correspond to expression at the endogenous *hb* loci and one of them corresponds to expression at the *hb-18kb* BAC locus. No expression is detected in the posterior. (**C-D**) Embryos are homozygous for the *5’MS2-hb-18kb* BAC insertion (**C**) or heterozygous for the *3’MS2-hb-18kb* BAC insertion (**D**). Top: Expression map of a typical nc11 embryo detected with the *hb* probe (green) and with the *MS2* probe (orange). As highlighted on one example (bottom left), nuclei in the anterior exhibit four sites of ongoing transcription detected with the *hb* probe and two sites detected with the *MS2* probe. Two of the *hb* sites co-localize with the two *MS2* sites and correspond to expression from the BAC. The two other *hb* site reveal ongoing transcription at the endogenous *hb* loci. In the posterior (bottom right), only two FISH signals are detected and co-localize with both the *hb* probe and the *MS2* probe. These two sites correspond to ongoing transcription at the BAC locus. **E)** Mean density of active nuclei in the posterior detected by the *hb* probe (green) or by the *MS2* probe (orange) on nc11 embryos carrying the *hb-18kb BAC (hb-18kb*), the *5’MS2-hb-18kb* BAC (*5’MS2*) or the *3’MS2-hb-18kb* BAC (*3’MS2*) insertion. Mean are calculated for n embryos and error bars correspond to standard deviation.

Taking advantage of BAC recombineering, we generated *MS2-hb-18kb* transgenes carrying insertions of the MS2 cassette either in the 5’UTR within the intron of *hb (5’MS2-hb-18kb*) or in the 3’UTR of *hb (3’MS2-hb-18kb).* Expression of these new MS2 reporters was assessed by live imaging (SI, Movie2) and double RNA FISH using a *hb* probe and a *MS2* probe. In the anterior of nc11 embryos either homozygous for the *5’MS2-hb-18kb* transgene (Fig 3C) or heterozygous for the *3’MS2-hb-18kb* transgene (Fig 3D), the *hb* probe highlights at most four spots revealing ongoing transcription at the two endogenous *hb* and two (C) or one (D) *MS2-hb-18kb* loci. In most cases, two of the *hb* spots co-localize with two (C) or one (D)*MS2* spot(s), which specifically label ongoing transcription at the *MS2-hb-18kb* loci. In these embryos, we also detect *hb* and *MS2* spots in posterior nuclei (Fig 3, C-D) which for most of them co-localize, thus revealing ongoing transcription at the *MS2-hb-18kb* loci. Posterior expression of the *MS2-hb-18kb* loci is also detected in living embryos expressing the MCP-GFP (Movie2). Altogether, these data strongly argue that posterior expression of the *MS2-hb-18kb* transgenes is mediated by the MS2 cassette and suggest that Zelda-dependent posterior expression of the *hb-MS2* reporter is mediated by cis-acting sequences in the MS2 cassette.

### The hb canonical promoter is sufficient to recapitulate early zygotic expression of hb

Given the trans-acting effect of Zelda on the posterior expression of the *hb-MS2* reporter (Fig 2) and the enhancer-like behavior of the MS2 cassette for posterior expression (Fig 3), we searched for potential Zelda binding sites in the MS2 sequence, using the ClusterDraw2 online algorithm (27) and Zelda position weight matrix (28). The canonical Zelda binding site is a heptameric motif (CAGGTAG, Fig 4A) over represented in the enhancers of pre-cellular blastoderm genes (23, 29). Strikingly, in the sequence of the MS2 cassette we find the motif CAGGT<u>C</u>G (a single mismatch with the canonical Zelda site) repeated 12 times and the motifs <u>T</u>AGGTA<u>C</u> (two mismatches) and <u>T</u>AGG<u>C</u>A<u>A</u> (three mismatches) each repeated 12 times (Fig 4B). This *in silico* analysis indicates that the MS2 sequence contained a total of 36 potential Zelda binding sites all located within linkers between MS2 loops (Fig 4B). Although we cannot be conclusive on the affinity strength of those various binding sites for Zelda, all of them share high similarity with TAGteam motifs (23). We thus engineered a new MS2 cassette mutating the 36 putative Zelda binding sites of our original *hb-MS2* reporter and inserted this new MS2-ΔZelda cassette under the control of the *hb* canonical promoter as in the original *hb-MS2* reporter. Expression of this new *hb-MS2 ΔZelda* reporter was assessed in living embryos expressing the MCP-GFP protein (SI, Movie3). Remarkably, we do not detect any MS2 spots in the posterior of the embryo (Fig 5A) indicating that unlike our original *hb-MS2* reporter (18), the new *hb-MS2 ΔZelda* reporter is expressed exclusively in the anterior at early cycles 10 to 11 as detected for endogenous *hunchback* by RNA FISH (3).

**Fig 4:**
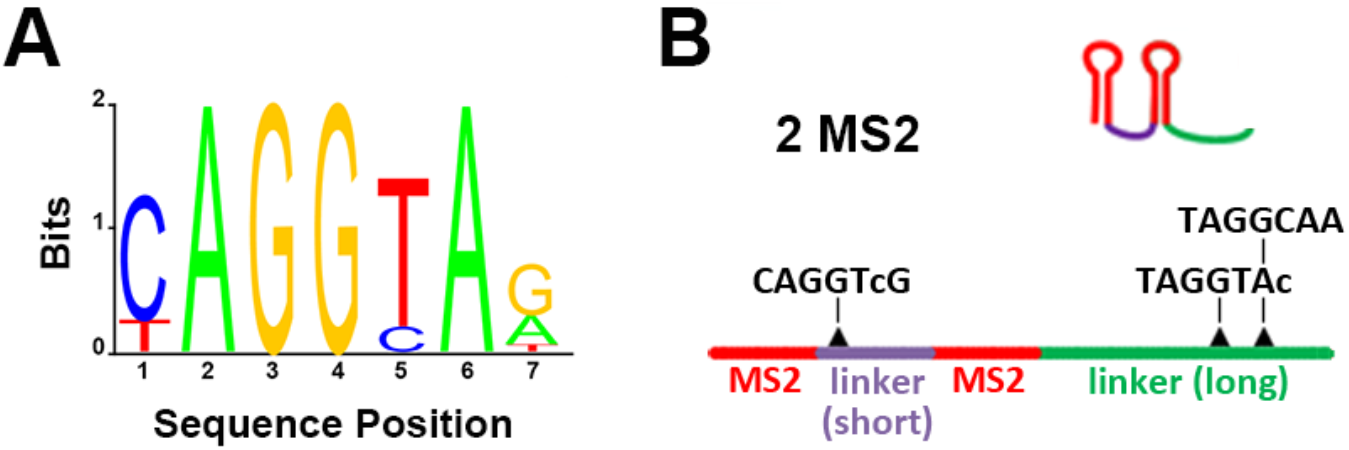
Zelda putative binding sites in the MS2 cassette: **A)** Position weight matrix of the Zelda binding site (28). The critical positions in the canonical Zelda site **CAGGTAG** are highlighted in bold. **B)** The 24 MS2 stem loop repeat corresponds to 12 tandem repetitions of the two MS2 repeats shown. In this sequence, the MS2 loops (red) are separated by either a short (purple) or a long (green) linker. The short linker contains one CAGGT<u>C</u>G sequence which harbors a single but critical mismatch with the canonical Zelda site. The long linker contains one <u>T</u>AGGTA<u>C</u> and one <u>T</u>AGG<u>C</u>A<u>A</u> which harbor respectively two or three permissive (not critical) mismatches with the canonical Zelda site.

**Fig 5:**
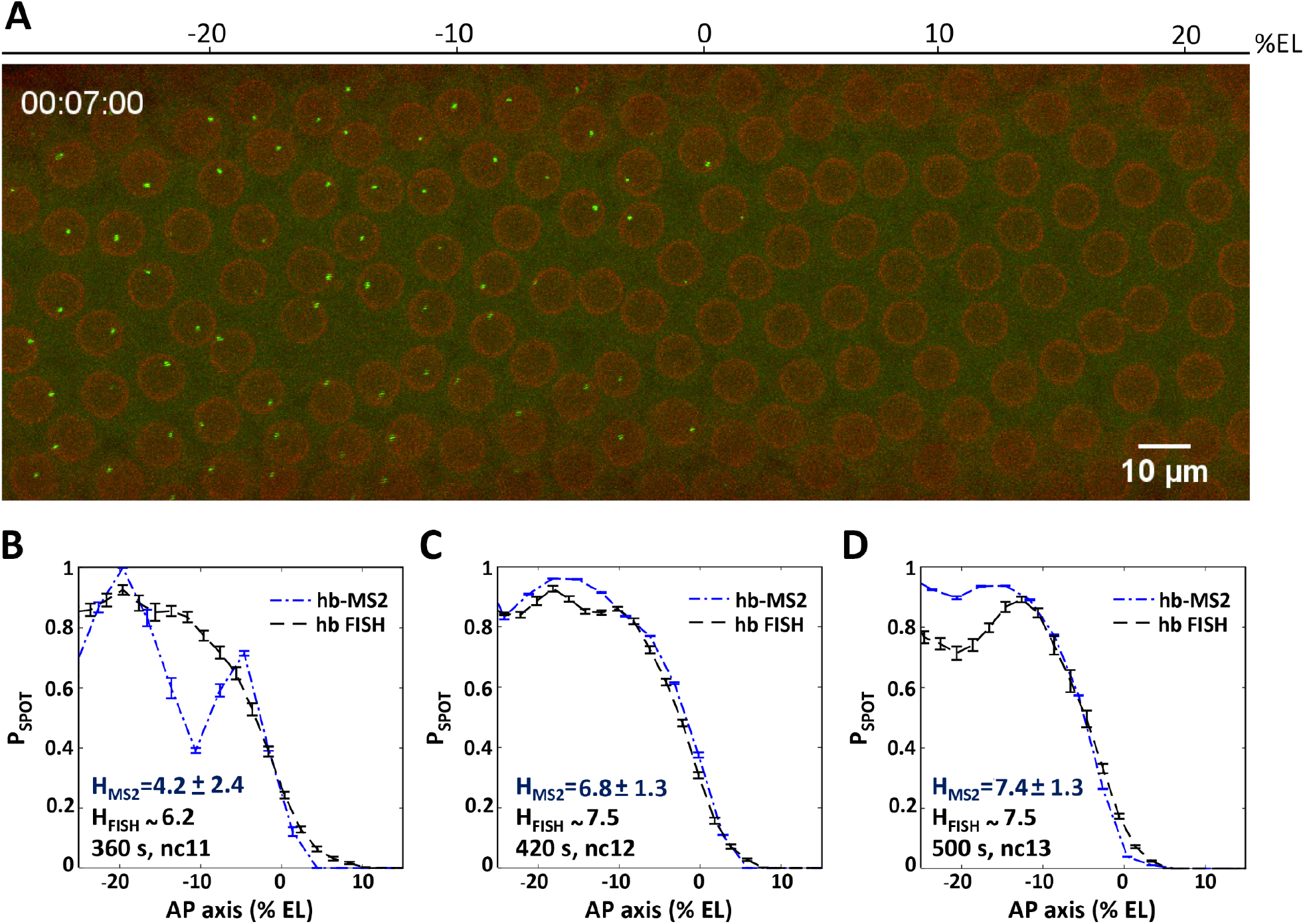
The *hb* canonical promoter expresses the *MS2ΔZelda* cassette in an anterior domain with a steep posterior boundary. (A) A 2D maximum projection snapshot from Movie3 (SI) was taken at ~7 minutes after the onset of nc12 interphase. In the green channel, MCP-GFP proteins recruited by the nascent MS2-containing mRNA can be seen accumulating at the *hb-MS2ΔZelda* loci (bright spots). In the red channel, the mRFP-Nup proteins localized at the nuclear envelope delineate nuclei. (B-D) Probability of active *hb-MS2 ΔZelda* loci along the AP axis (dashed blue lines with error bars), extracted from snapshots of 6 movies near the end of each nuclear cycle: 360 s after the onset of nc11 interphase (B), 420 s after the onset of nc12 interphase (C) and 500 s after the onset of nc13 interphase (D). In each panel the probability of active endogenous *hb* loci (black dashed lines with error bars) extracted from the FISH data from (3) is also shown. Hill coefficients (H) are indicated in blue (*hb-MS2* reporter) or in black (endogenous *hb* from FISH data).

As RNA FISH are performed on fixed embryos whereas the MS2 data are obtained from live material, we wondered whether these two different approaches provide consistent quantification of *hb* transcriptional activity when focusing on the “steepness” of the expression boundary. Therefore, the probability to be active for *hb-MS2ΔZelda* loci at a given position along the AP axis and at a given time (Pspot) was extracted from movie snapshots and compared to RNA FISH data of endogenous *hb* expression (3). To compare data from several embryos, embryos were aligned fixing the origin of the AP axis when the probability for a locus to experience transcription at any time during the interphase (Pon) is equal to 0.5 (for the definition of the boundary see details in SI, Section IV). This allows us compensating for input noise (noise in the Bicoid gradient) which was shown to be of about 2-3 *%* EL (6) and focusing only on the output noise (noise in the transcriptional response). As shown in Fig 5 (B-D), the curves plotting the mean spot appearance P_SPOT_ as a function of position along the AP axis are remarkably similar when extracted either from the RNA FISH data (dashed line, (3)) or the MS2 movie snapshots (blue lines) with Hill coefficients varying from 4.5 (nc11) to ~ 7 at nc12 and nc13. Thus, the *hb-MS2 ΔZelda* reporter is expressed at early cycles in an anterior domain with a boundary as steep as the boundary of the endogenous *hb* expression domain. These experiments show that the *hb* canonical promoter is sufficient to faithfully recapitulate the early zygotic expression of endogenous *hb.* This is in contradiction with our first interpretation (18) deduced from the expression of the original *hb-MS2* misleading reporter with its high number of unexpected Zelda binding sites. Nevertheless, the now almost perfect match between Pspot activity along the AP axis from the FISH data of endogenous *hb* expression at the end of the interphase and data extracted from movies of the *hb-MS2ΔZelda* reporter, ensures that the analysis of this reporter expression dynamics can help understand how the step-like expression of *hb* arises.

### The hb-MS2ΔZelda reporter shows different transcription kinetics in the anterior domain

The MS2 movies provide access to the transcription dynamics of each *hb-MS2 ΔZelda* single locus in all the visible nuclei of developing embryos. Key features are extracted from the time traces of the MS2-GFP spots (Fig 6A), choosing as the origin for time the onset of interphase for that particular nucleus (see details in SI, Section III). These features include: *i*) the initiation time (t_init_) which measures the time period it takes after the onset of interphase to detect the first MS2 transcription signal at the locus, *ii*) the time period during which the locus is activated (t_active_) and *iii*) the time period at the end of interphase during which the locus is turned off (t_end_). From the time traces (Fig 6A), the integral activity (ΣI) integrates the area under the trace and provides a relative measure of the total amount of mRNA produced. The average mRNA production rate (μI) is calculated by dividing ΣI by t_active_. Features are obtained from 5 (nc11), 8 (nc12) and 4 (nc13) embryos. Embryos were aligned spatially fixing the origin of the axis at boundary position (P_ON_) at nc12 and the origin of time was calculated for each nuclei as the origin of its respective cycle (see SI, Section III). As shown in Fig 6, several of these features exhibit different behaviors depending of the position along the AP axis. Notably, t_init_ and ΣI appear more variable when loci are located close to the boundary than in the most anterior part (Fig 6B-G). Similarly, in a region of about 10 % EL at the boundary, the mRNA production rate drops from the constant value reached in the anterior region to 0 at the boundary (Fig 6H-J). Thus, the dynamics of the transcription process at the *hb-MS2ΔZelda* reporter exhibit two distinct behaviors: in the anterior of the embryo, time trace features are stable likely reflecting a maximum PolII loading rate and saturated levels of Bicoid; in a region of ~ 10 % EL anterior to the expression boundary, time trace features are fluctuating reflecting limiting amounts of Bicoid. The pattern steepness corresponds to a Hill coefficient of ~7 to 8 of the regulation function (see SI section IV). Tailor-made analysis of the time traces allowed us to extract different kinetic parameters of promoter activity in these two regions and to demonstrate that transcription of the *hb-MS2ΔZelda* reporter is bursty (described by a two states model) in both regions (30). Thus, MS2 live imaging gives us the opportunity to really decipher the dynamics of the *hb* expression compare to the FISH that was just giving us P_SPOT_ (probability for *hb-MS2ΔZelda* loci to be active at a given position along the AP axis and at a given time).

**Fig 6:**
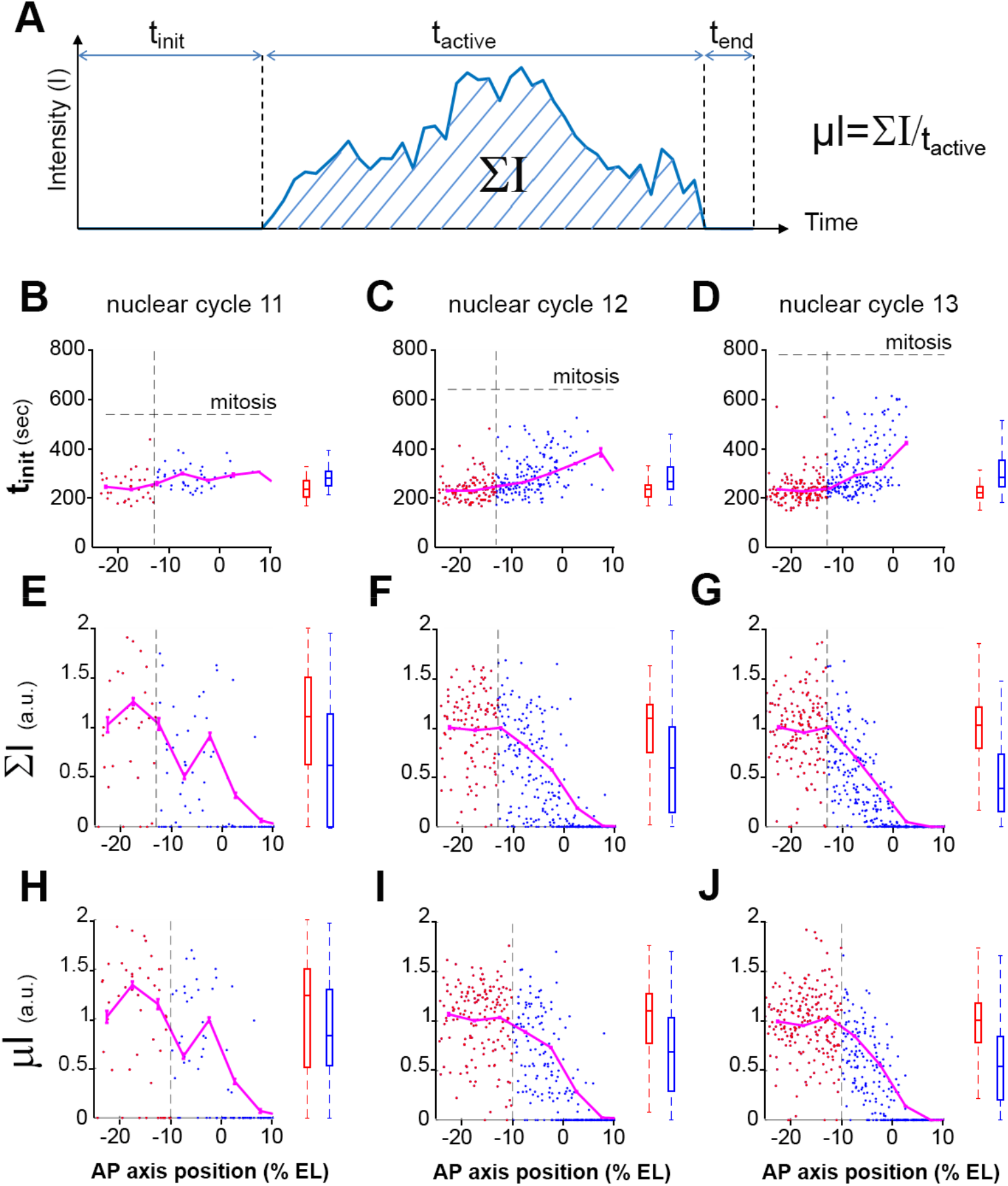
Distribution of time traces features along AP axis. (A) Description of the time traces features. Initiation time (t_init_, B-D), normalized integral spot intensity (ΣI, E-G) and production rate (μl, H-J) are indicated as a function of position along the AP axis with the origin fixed for each embryo at the position of the expression boundary at nc12. Embryos were at nc11 (B, E and H), nc12 (C, F and I) and nc13 (D, G and J). The vertical dashed line indicates when the significant change in the feature distribution from the anterior pole is first detected in nc13, using KS test (p-value 0.05). This line separates the *hb* expression domain into 2 zones: the anterior zone which exhibits a saturating process (red box) and the boundary zone in which exhibit a more stochastic activity characterized by more variability (blue box). On the right of each panel (B-J), the distribution of the two populations is shown (median + 25% and min/max). Example of MS2-MCP time traces are given in SI (Section II). Data were obtained from 5 (nc11), 8 (nc12) and 4 (nc13) embryos. Embryos were aligned spatially fixing the origin of the axis at boundary position (P_ON_) at nc12 and the origin of time was calculated for each nuclei as the origin of its respective cycle (see SI, Section III). For t_init_, the data points are for expressing nuclei only whereas for ΣI and μl both expressing and non-expressing nuclei were taken into account.

### At each nuclear cycle, the hb expression pattern reaches steady-state in 3 minutes

The temporal dynamics of the *hb* pattern establishment at the scale of the whole embryo (along the AP axis) were extracted from the MS2 movies: the probability for the locus to be ON as a function of the position along the AP axis and time in the cycle (P_SPOT_) can be visualized in Movie4 (SI) and is plotted in the form of kymographs on pulled embryos as above (Fig 7A-C). At each cycle, spots can appear as early as ~ 150 s after mitosis. Given the interruption of transcription during mitosis, this limit of 150 s corresponds to the period required to re-establish transcription during the interphase and likely includes genome de-condensation, the time it takes for the Bicoid protein to be imported in the nucleus after mitosis and the time it takes for the MS2 system to produce a signal that is above background. After this period, expression rapidly turns on in the anterior to reach the plateau value where all nuclei express the reporter (P_SPOT_ ~ 1). The time period to reach the plateau is shorter at the most anterior position in the field of view imposed by the movie recording (~ – 25% EL) and increases with the distance to the anterior pole (Fig 7D-F) consistent with a dose-dependent activation process of *hb* by Bicoid (4, 8). Then, the expression pattern is steadily established, which includes fixing the boundary position and steepness. This stable state lasts for a period of time which varies with the length of the cycle until a rapidly gets switch off simultaneously along the AP axis: at 400 s at nc11 (Fig 7, A & G), 500 s at nc12 (Fig 7B & 7G) and 750 s at nc13 (Fig 7C & 7G). Importantly, the dynamics of the boundary positioning is the same at the three nuclear cycles considered (Fig 7G) and the steady state of boundary positioning is reached rapidly: ~330 s after the onset of the cycle corresponding to ~ 180 s after the first hints of transcription at this position (Fig 7G). Dynamics of pattern steepness also exhibit similar behaviors (see SI, section VI). Thus, expression of the *hb-MS2ΔZelda* reporter allows us to directly observe position-dependent transcriptional activation which is consistent with Bicoid dose-dependent transcriptional activation. Also, it demonstrates that the steady-state of positional measurement is reached in no more than 3 min at each cycle with very similar dynamics between cycles.

**Fig 7:**
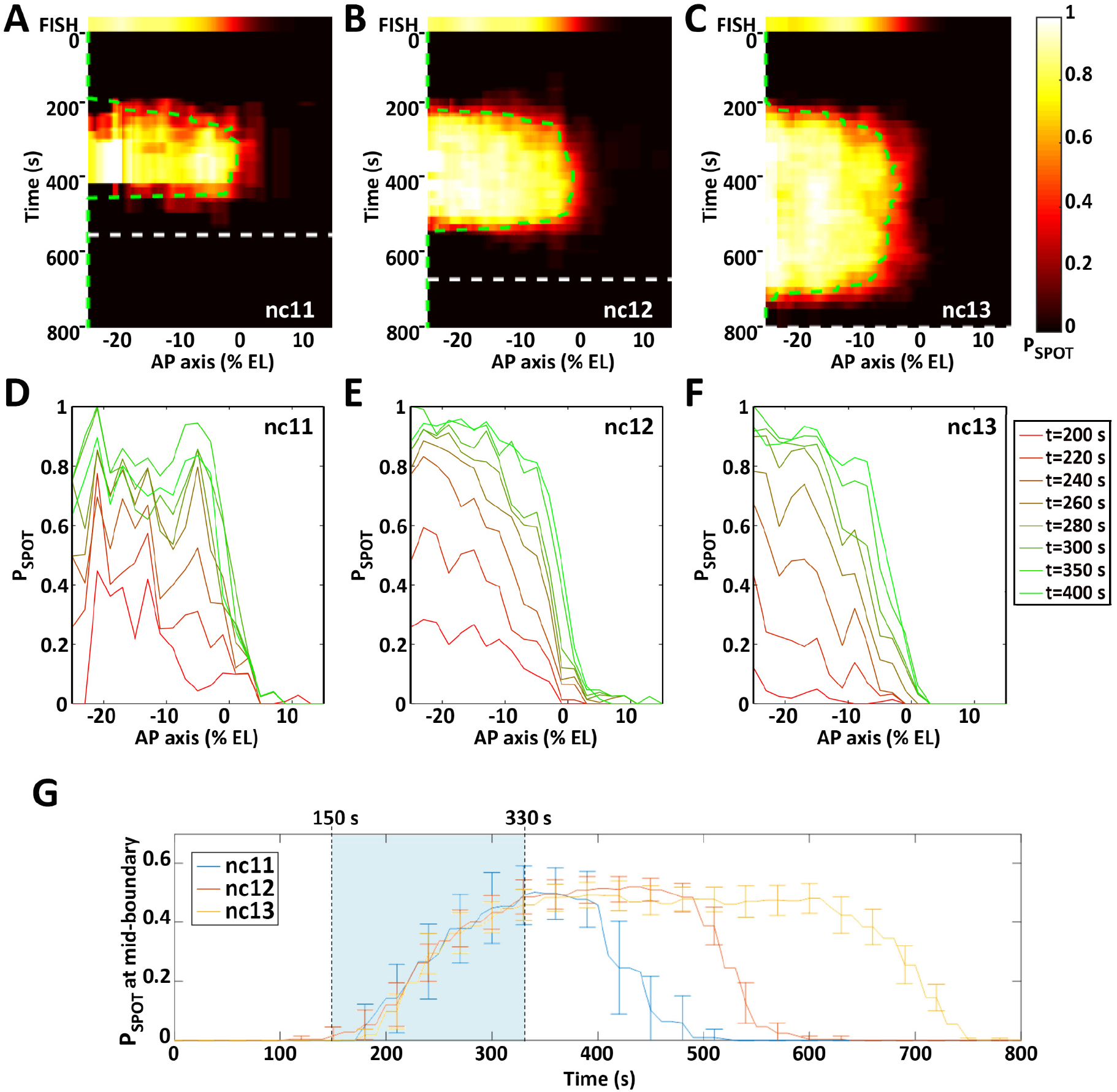
The dynamics of *hb-MS2ÁZelda* expression pattern. **A-C**: The probability for a given locus to be ON (P_Spot_) is indicated by a heat map (color scale on the right where P_Spot_ = 0 is black and P_spot_ = 1 is white) horizontally as a function of position along the AP axis (0% EL positioned Pon boundary at nc12) and vertically as a function of time (s) fixing the origin at the onset of interphase for each nucleus (see details in SI, Section III). On the top the endogenous *hb* expression pattern as measured from RNA FISH (16) are shown. For each cycle (**A**: nc11; **B**: nc12; **C**: nc13), the end of interphase (onset of next mitosis) is indicated by a dashed line (white). The green dashed line indicate the position of the expression boundary (P_Spot_ = 0.5) over time. **D-F**: Pspot as a function of position along the AP axis for different time during the interphase of nc11 (**D**), nc12 (**E**) and nc13 (**F**). **G**: P_Spot_ as a function of time (s) at mid-boundary position (where P_Spot_ reach a steady value of 0.5). The first hints of transcription are observed at mid-boundary position ~ 150 s after the onset of interphase (lower limit of the light blue zone) and steady state is reached at ~ 330 s (higher limit of the light blue zone). Boundary formation reaches steady state in ~ 180 s. Data were obtained from 5 (nc11), 8 (nc12) and 4 (nc13) embryos. Embryos were aligned spatially fixing the origin of the axis at boundary position (Pon) at nc12 and the origin of time was calculated for each nuclei as the origin of its respective cycle (see SI, Section III).

### Modeling the dynamics of the hb expression pattern

Surprisingly, the *hb* boundary is fixed at a given position early during the interphase. To better understand how the stable response downstream of Bicoid is achieved, we build a stochastic model of *hb* expression regulation by the Bicoid transcription factor (TF), coupled with a stochastic transcription initiation process assuming random arrival of RNA polymerases when the gene is activated (Fig 8A). The mechanism of *hb* expression regulation through the cooperative binding of multiple TFs to the promoter was originally proposed in 1989 to explain how the shallow Bicoid gradient could give rise to an expression pattern with a steep boundary (8, 9). It was subsequently proposed (31) that within an equilibrium binding model, a pattern steepness quantified by a Hill coefficient of *N* requires a promoter with at least *N* TF binding sites. Thus, we modelled *hb* expression regulation through the binding and unbinding of TFs to *N* identical operator sites on the *hb* promoter (see Fig 8A and Methods section). Here, TF concentration at the boundary position is rate-limiting as it takes time for a diffusing TF to reach binding sites at the promoter (see SI section VII.1).

**Fig 8:**
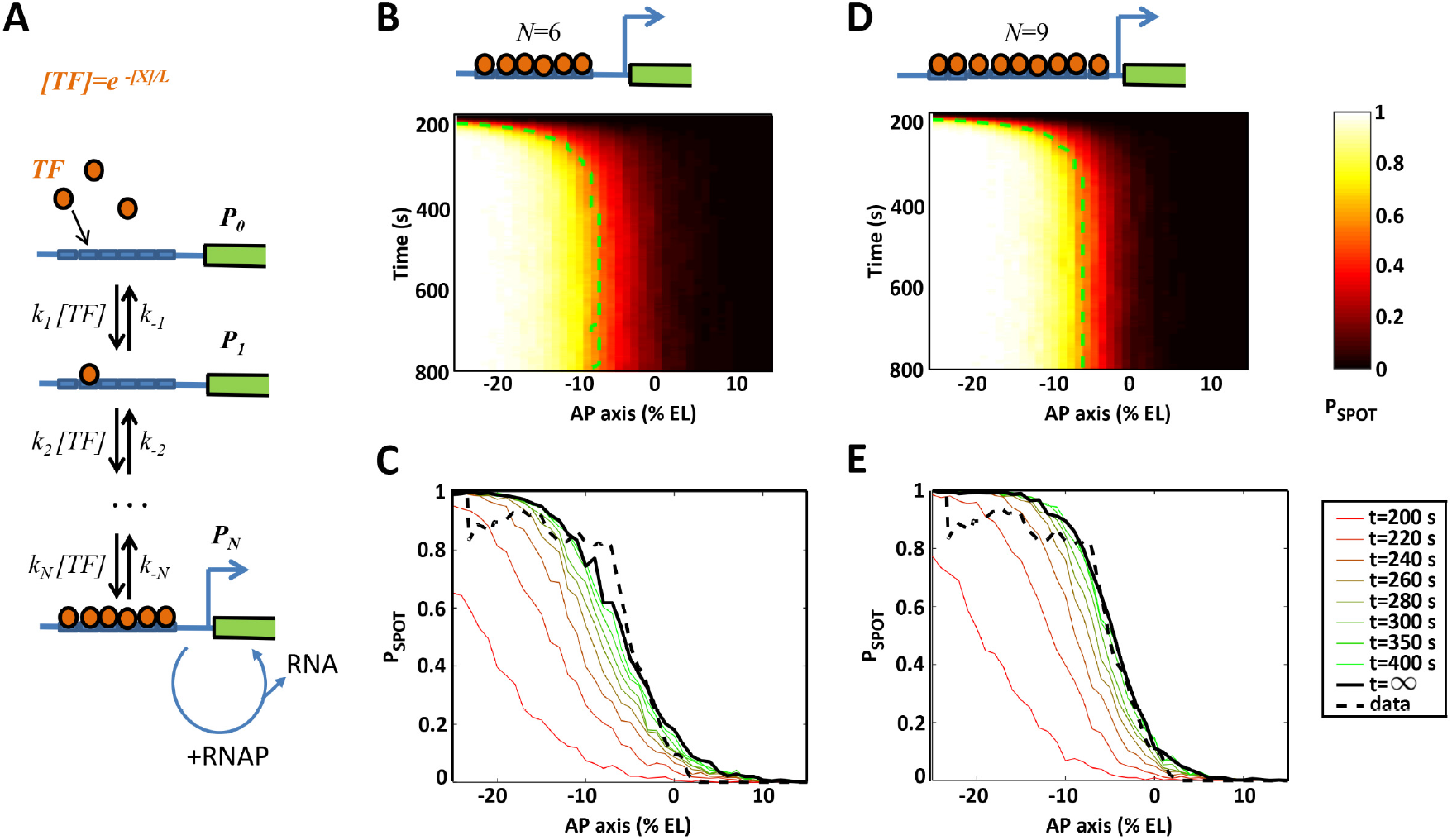
Modeling transcriptional regulation by the Bicoid transcription factor through interactions with the *hb* promoter operator sites. **A)** Model of regulation by the Bicoid transcription factor (TF) binding to multiple binding sites on the *hb* promoter coupled with stochastic transcription initiation. Transcription initiation is allowed only when the binding sites are fully bound. During this window, RNAP can randomly bind to the promoter and initiate transcription to produce mRNA. **B** or **D**) The model prediction for the probability of an active transcription locus (P_SPOT_, colorbar) as a function of time in the nuclear cycle and position along the AP axis for a model with 6 (**B**) or 9 (**D**) Bicoid binding sites. **C** or **E**) The simulated pattern evolution of P_SPOT_ along the AP axis over time (colored line), shown with the pattern predicted at steady state (solid black line) and the stable pattern extracted from the data in nc13 (dashed black line, as in Fig 7F). Panels C and E represent cuts in time of panels **B** and **D**, respectively. The kinetic parameters were chosen so as to match *hb* pattern’s steepness and formation time at the boundary (P_SPOT_=0.5) (see Table S3). The value of P_SPOT_ is calculated from 200 trajectories per AP position.

Motivated by the 6 known Bcd binding sites on the *hb* promoter (8, 32), we first consider a model with 6 binding sites (N=6). The binding and unbinding rates are chosen (see SI section VII.5) to achieve the closest steepness and establishment period from the *hb* pattern observed from the movies (Fig 7. A-C). The model fits are performed on data pulled from all embryos and nuclear cycles. The gene expression pattern dynamics, shown as the probability (P_SPOT_) of a nucleus having an active locus (bright spot) as a function of position along the AP axis and as a function of time during the nuclear cycle, shows a good qualitative agreement between the model and the data. In Fig 8B, similarly to Fig 7A-C, initially there is no expression since the TF are not bound to the promoter. Active transcription loci first appear near the anterior pole, where the activator concentration is the highest, and then transcription activation propagates quickly to the mid-anterior region. After a certain amount of time, the steep expression pattern becomes stable, with the boundary (dashed green line) located near the mid-embryo region where the *hb* boundary is located. However, using the canonical number of binding sites (N=6) results in a steepness of the boundary lower than that observed in the experimental data (Fig 8C). Increasing the binding sites number from 6 consecutively to 7, 8, 9 or 10 allows getting closer to the experimental data, with the observed boundary steepness recovered assuming *N=9* binding sites (Fig 8E). Increasing *N* beyond 9 did not result in significantly better fit (see SI section VII.5). On the other hand, due to the trade-off between the pattern steepness and the promoter switching rate [32], accommodating the observed steepness using *N* slightly larger than the observed Hill coefficient (~7) results in a very long formation time. Thus, within the constraints of our model, it seems that there is no way to account both for the rapid establishment and the steepness of the transcriptional response.

## Discussion

In this study, the removal of 36 putative binding sites for the transcription factor Zelda (unfortunately present in the sequence of the MS2 cassette) reveals a new temporal dynamics of the *hb* canonical promoter at the onset of zygotic transcription. Importantly, unlike our original *hb-MS2* reporter (18), the new *hb-MS2ΔZelda* reporter faithfully reproduces the early zygotic expression of the endogenous *hb* observed with RNA FISH (3, 18). It indicates that the 750 bp of the *hb* locus, including 300 pb of the proximal enhancer, the zygotic promoter and the intron, are sufficient to reproduce the endogenous expression of the *hb* gene in the early nuclear cycles (11 to 13). The dynamics of establishment of the *hb* pattern is thus properly captured by the new *hb-MS2 ΔZelda* reporter.

These MS2 movies provide access to the *hb* pattern dynamics, which was not perceivable in previous *in situ* experiments on fixed embryos. *hb* expression first occurs in the anterior then proceeds to the boundary region. A difference in the activation time (initiation time) following mitosis is observed even within the anterior region, allowing us to visualize for the first time position-dependent activation of *hb* and thus likely dose-dependent activation by Bicoid. Analysis of the MS2 time traces indicates that the transcription process is more variable among nuclei at the boundary of the expression domain, where Bicoid concentration is low and probably limiting, than in the anterior where the concentration of Bicoid is high. This anti-correlation between the relative variability of mRNA production among nuclei and the Bicoid concentration (position along the AP axis) supports the idea that Bicoid interactions with the *hb* promoter are rate-limiting processes contributing to “bursty” transcription (30). Nevertheless, rate-limiting interaction of Bicoid with DNA is not the sole factor contributing to bursty transcription as it is also observed in the anterior region with very high Bicoid concentration (30). Thus, despite an extremely fast transcription initiation imposed by the 5 min interphase and frequent mitoses, bursty transcription is clearly observed at the onset of zygotic transcription in fly embryos.

The MS2 movies indicate that the *hb* boundary is established within 3 minutes at each nuclear cycle with a very high steepness (H ~7). How the steepness and positioning of the boundary are reached so rapidly is unclear. Given the different transcription features observed in the anterior vs boundary, Bicoid is likely to be a rate limiting factor in the formation of the *hb* pattern around the boundary region. Therefore, the interactions between Bicoid molecules and the *hb* promoter need to be modeled explicitly, rather than implicitly. We propose a model of transcription regulation via the binding/unbinding of transcription factors to the operator sites of the target promoter. Using this model, we show in a tandem paper that accommodating a steep gene expression pattern requires very slow promoter dynamics and thus very long pattern formation time (33). Considering this tradeoff, the steepness and the pattern formation time observed from the movies are even more intriguing. The most relevant model with 6 binding sites, motivated by previous works (8), fits well the pattern dynamics but fails to reproduce the observed steepness of the boundary in such a short time period. The failure of the 6 binding sites model to completely reproduce the experimental data indicates that additional mechanisms are required to enhance the steepness of the boundary to the observed level. It was recently proposed that energy expenditure, encoded as non-equilibrium binding of the TF, can allow a Hill coefficient greater than the number of binding sites (up to 11 with 6 sites) (21). Alternatively, different regulatory scenarios could play a role. First, we found that increasing the number of binding sites up to 9 allows a significantly better fit of the model with the data than with 6 binding sites. It is thus possible that the 750 bp of the *hb* gene that are sufficient to elicit the pattern contain more than 6 Bicoid binding sites. As recently pointed out, the importance of low affinity binding sites might be critical to confer specificity and robustness in expression (34) and a closer analysis of the *hb* regulatory sequence looking for potential weak binding sites for Bicoid might clarify this point. A second possibility is the involvement of other transcription factors distributed as gradients that could bind to the *hb* promoter and contribute to the increase in the steepness of the boundary. The Hb maternal protein, which is also expressed as an anterior to posterior gradient and able to bind to the endogenous *hb* promoter (35), contributes to the *hb* expression process by allowing expression at lower Bicoid concentration thresholds (12) and faster activation (16). Although this has not yet been investigated, maternal Hb might also contribute to sharpen the *hb* boundary in a timely manner. Alternatively, maternal repressors expressed as gradient in the posterior region or downstream of the boundary could also contribute to the steepness of the *hb* boundary. Among potential candidates are Caudal, which is expressed as a posterior to anterior gradient but so far has been described as a transcriptional activator in fly embryos (36), or Capicua, a transcriptional repressor at work in the center of the AP axis, where the *hb* boundary forms (7).

From the movies, the dynamics of establishment (before reaching steady state) of the *hb* pattern appears to be invariant at the three nuclear cycles considered (nc11, nc12 and nc13). At these three nuclear cycles, it takes ~180 s from the detection of the first MS2-MCP spots to the establishment of the pattern near mid-embryo position (Fig 7G). This invariance in the dynamics of establishment indicates that there are no dramatic changes in the regulation of the transcription process during the three cycles suggesting that Bicoid remains one of the main patterning factors of *hb* transcription dynamics at the boundary region in these stages of development. The other transcription factors involved in *hb* expression (16, 37), if any, would need to be stably maintained during the three cycles. As mentioned above, one possible factor is the Hb protein itself which was shown to contribute to the expression of the *hb* expression pattern with both a maternal and a zygotic contribution (12, 16). A likely hypothesis is that up to nc13, the zygotic Hb protein production is balanced with the maternal Hb protein degradation, thus leading to a stabilized Hb gradient over the three considered cycles.

Previous observations of Bicoid diffusion in the nucleus space pointed out that it requires at least 25 minutes for a single Bicoid binding site to sense the Bicoid concentration with 10% accuracy, a level observed using FISH and protein staining experiments (5, 16, 38). This estimation is based on the Berg-Purcell limit in the precision of concentration sensing of diffusing molecules via surface receptors (39–41). It should be noted that the Berg-Purcell limit applies when the receptors’ interactions with TF are independent. In such a situation regarding the Bicoid/*hb* system, the resulted gene expression pattern is non-steep (*H* ~ 1.31, see SI Table S2). In-depth analysis of the model in the tandem paper (33) shows that increasing the pattern steepness slows down the switching rate between the gene’s active and inactive states, due to the required cooperativity between the binding sites (21). Consequently, our fitted models (with *N=6* or *N=9* binding sites) result in much higher errors in the integrated readout when compared to the model of independent binding sites (SI Fig S10), and require a significantly longer integration time to achieve a specific precision level. Thus, the answer to the precision of the *hb* pattern achieved in such a short interphase duration remains elusive. In the future, systematic studies using synthetic promoters with a varying number of Bicoid binding sites and quantitative analyses of promoters’ dynamics captured with MS2-MCP system will help characterize not only the cooperativity of Bicoid binding sites but also the kinetics of the downstream processes once the gene is activated. Added to this, the Bicoid search time for its binding sites on *hb* (16, 38) also needs to be revisited. The employed value (~4 s) in our model is estimated assuming a 3D search process inside the nucleus space but if Bicoid can slide along DNA in search for the sites, the process can be ~100 times faster (42). Such a possibility is compatible with fluorescence correlation spectroscopy measurements of Bcd-eGFP motion (16, 43) and the recent observed clustering of the Bicoid molecules across the embryo (44). The advent of single molecule tracking methods (44, 45) represent a promising approach to further shed lights in the mechanism of such process.

Importantly, despite these very rapid and precise measurements of Bicoid concentration along the AP axis, the expression process itself shows a great variability in the total amount of mRNA produced during the interphase per nucleus (δmRNA/<mRNA> ~150% in the boundary region) (30). It is thus difficult to gauge to which degree errors in sensing Bicoid concentration contributes to the variability of the total mRNA produced at the boundary (30). This also raises the question of understanding how precision in the downstream processes required for embryo segmentation is achieved at the scale of the whole embryo. If the embryo is capable of spatially averaging *hb* expression between nuclei at the same AP position (for example by the diffusion and nuclear export of its mRNA and nuclear import of its protein) (46), this will help the system distinguishing between the anterior and posterior region based on the *hb* transcription pattern alone. However, spatial averaging has a limit, due to the very short time available, the limited number of nuclei and the finite diffusion coefficient of the *hb* gene’s products (46). It is likely that nuclei need to integrate *hb* gene expression over the nuclear cycle to reduce the noise in the *hb* readout. In nc11, the pattern collapses soon after reaching steady-state due to the next mitosis round. Therefore, any integrations of gene expression in nc11 and earlier (interphase duration shorter than ~400 s) are likely to lead to bias the pattern boundary to the anterior. Only starting at nc12 is the interphase duration long enough to reliably observe a steep pattern with the border around mid-boundary position, based on the number of mRNA produced per nucleus (ΣI).

## IV. Materials and Methods

### Drosophila stocks

The original reporter *hb-MS2* and the Nup-RFP and MCP-GFP transgenes, both inserted on the second chromosome, were from (18). All transgenic stocks generated in this study were obtained by BestGene. The *zld^294^FRT19A* chromosome was a gift from C. Rushlow (23) and female germline clones were induced using heat-shock FLP recombinaison (47) with a *OvoD1hsFLPFRT19A* chromosome (# 23880, Bloomington). All stocks were raised at 25°C.

### Plasmids and BACs

The *hb-18kb-BAC* spanning the *hb* locus was the BAC CH322 55J23 obtained from PACMAN BAC libraries (48). The *24xMS2 cassette* was inserted in the 5’UTR within the *hb* intron (*5’MS2-18kb-BAC*) or in the 3’UTR (*3’MS2-18kb-BAC*) using BAC recombineering (49) (details in SI-Section I). The plasmid used to generate the new *hb-MS2ΔZelda* reporter was generated by replacing in the *pCasPeR4-hb-MS2* construct from (18), the MS2 cassette by a new MS2 sequence synthetically generated by Genescript in which all putative Zelda binding sites (*24xMS2-SL-ΔZelda*) had been mutated. The sequence coding for the CFP in the *pCasPeR4-hb-MS2* construct from (18) was also replaced by the sequence coding the iRFP (Addgene 45457) in which a unique putative Zelda binding site had been mutated (See SI-Section I for more information).

### The MS2-SL-ΔZelda Sequence

Sequence-based binding-site cluster analysis was performed using the online software ClusterDraw2 v2.55 at line.bioinfolab.net/webgate/submit.cgi. PWM for Zelda has been generated from (28) using the following Zelda binding sites: CAGGTAG; CAGGTAA; TAGGTAG; CAGGTAC; CAGGTAT; TAGGTAA; CAGGCAG and CAGGCAA. All heptamers detected as being a potential Zelda binding sites have been mutated and the new sequence synthesized by Genscript. The sequence of the new MS2 ΔZelda cassette is given in SI (Section I).

### RNA FISH and analysis

Embryo fixation and RNA *in-situ* hybridization were performed as describe in (3). Briefly, RNA probes were generated using T7/T3 *in-vitro* transcription kit (Roche). *hb* RNAs were labelled with digoxigenin-tagged anti-sense probes detected with a sheep anti-dig primary antibody (1/1000 dilution, Roche) and donkey anti-sheep Alexa 568 secondary antibody (1/400 dilution, Invitrogen). MS2 containing RNAs were labelled with biotin-tagged anti-sense probes detected with a mouse anti-bio primary antibody (1/400 dilution, Roche) and chicken anti-mouse Alexa 488 secondary antibody (1/400 dilution, Invitrogen). Embryos were incubated 10 min with DAPI for DNA staining and 10 min in WGA-Alexa 633 (1/500 dilution, Molecular Probes) for nuclear envelop staining. Fixed embryos were mounted in VectaShield (Vector) and imaged in 3D (~20Z x 0.45μm) with a XY resolution of 3040*3040, 8bits per pixels, 0.09μm pixel size, 1 airy unit using a Zeiss LSM780 confocal microscope with a Zeiss 40x (1.4 NA) A-Plan objective. A full embryo 3D RAW image is composed on three 3D RAW Images that were stitched using Stitch Image Grid plugins from FIJI with 10% overlap and the linear blending fusion method. Image processing and analyzing were performed as describe (18). Embryos of the proper stage in the nc11 interphase were selected according a threshold based on the nuclear area above 80 μm^2^ (corresponding to late interphase) as in (3). The expression map of endogenous *hb* or *MS2* transgenes have been manually false colored on FIJI and flatten on the nuclear channel.

### Live embryo imaging

Imaging conditions were comparable to those outlined in (18, 50). Embryos were collected 1h after egg laying, dechorionated by hand, fixed on the cover slip using heptane-dissolved glue, and immersed in to halocarbon oil (VWR). Mounted embryos were imaged at ~ 25°C on a Zeiss LSM780 confocal microscope with a Zeiss 40x (1.4 NA) A-Plan objective. Image stacks of the lower cortical region of the embryo close to the middle of the AP axis (pixel size 0.2 μm, 0.54 μs pixel dwell time, 8 bits per pixels, confocal pinhole diameter 92 μm, distance between consecutive images in the stack 0.5 μm, ~1200x355 pxl, ~30 z-stacks) were collected continuously. The GFP and RFP proteins were excited with a small fraction of the power output of a 488nm and a 568nm laser, 1.2% and 2% respectively. Images were acquired using the ZEN software (Zeiss). For each embryo, a tiled image of the the midsection of the whole embryo was obtained, by stitching 3 separate images, from which the position of the anterior and posterior poles could be inferred.

### Image processing and data extraction

Live imaging processing was performed in MATLAB as in (18). Following imaging, movies are checked manually to verify all the nuclei included in data analysis are fully imaged in their depth and incompletely imaged nuclei (mostly nuclei at the periphery of the imaging field) are excluded. Nuclei segmentation is performed in a semi-automatic manner using our own software (51). Only nuclei that exist throughout the nuclear interphase are used for the analysis. The MS2 spot detection was performed in 3D using a thresholding method. An average filter was applied before thresholding on each Z of the processed time point for noise reduction. MS2 spots were detected by applying a threshold equal to ~2 fold above background signal and only the spots composed of at least 10 connected voxels were retained. The intensity of the 3D spot is calculated as the sum of the voxel values of each Z-stack. At the end of nc13, some MCP aggregates may cause false spot detections: they are less bright and their shape is more spread compare to the MS2-MCP spots. The aggregates are eliminated automatically by raising the spot detection threshold the without affecting the detection of the MS2 spots. We manually checked each movie to ensure the correct spot detection. The data from a segmented movie indicates for each nucleus its segmentation profile, identifier number and the intensity trace of the detected spot over time.

During mitosis, nuclei divide in waves, usually from the embryo poles. Therefore, nuclei at the anterior pole may produce MCP-MS2 spots earlier due to either earlier chromatin decondensation or earlier reentrance of Bicoid into the nucleic space (6). We correct for this by realigning all the intensity traces by choosing the origin for time for each trace when the two sibling nuclei are first separated (see SI, Section III)

### Stochastic model of *hb* expression regulation

The general model of transcription regulation through transcription factor (TF) binding/unbinding to the operator sites (OS) is a based on a graph-based linear framework (21, 52, 53). We introduce a single time scale, which is the TF searching time for a single operator site at the boundary t_bind_ ~ 4s. The details of the model are described in SI (Section VII).

## V. Acknowledgments

The authors thank Patricia Le Baccon and the Imaging Facility PICT-IBiSA of the Institut Curie, BestGene Inc for transgenics and C. Rushlow and the Bloomington stock center for fly stocks. This work was supported by a PSL IDEX REFLEX Grant for Mesoscopic Biology (ND, AMW, MC), ANR-11-BSV2-0024 Axomorph (ND and AMW), ARC PJA20151203341 (ND), ANR-11-LABX-0044 DEEP Labex (ND), an Ontario Trillium Scholarship for International Students (CAPR), a Mitacs Global Link Scholarship (CAPR) and an Internal Curie Institute Scholarship (CAPR), a Mayent Rothschild sabbatical Grant from the Curie Institute (CF) and an NSERC discovery grant RGPIN/06362-15 (CF), a Marie Curie MCCIG grant No. 303561 (AMW) and PSL ANR-10-IDEX-0001-02. The funders had no role in study design, data collection and analysis, decision to publish, or preparation of the manuscript.

## VII. Supporting Information

### Movie S1. Live imaging of transcription dynamics of *hb-MS2 (Lucas et al 2013*) subject to varying dose of Zelda

(A) Zelda hetero (Zld Mat +/-): expression of the *hb-MS2* reporter in embryos from *zld^294^* heterozygous females. (B) Zelda GLC (Zld Mat -/-): expression of the *hb-MS2* reporter in *zld*^294^germline clone embryos. The movies have two channels: MCP-GFP channel (green) for monitoring the dynamics of nascent mRNA production and NUP-RFP (red) for nuclei detection. The capture frame is from −25% to 25% of embryo length. The anterior pole is on the left side of the frame. http://xfer.curie.fr/get/9uOHMTi46j0/mov1.avi

### Movie S2. Live imaging of the transcription dynamics with MS2 cassette inserted at 5’-UTR and 3’-UTR

MS2 cassette (*Lucas et al. 2013*) is placed at (A) 5 ‘-UTR within the intron of *hb* gene (*5’MS2-hb-18kb*) and (B) 3’-UTR of *hb* gene (*3’MS2-hb-18kb).* The movies have two channels: MCP-GFP channel (green) for monitoring the dynamics of nascent mRNA production and NUP-RFP (red) for nuclei detection. The capture frame is from −25% to 25% of embryo length. The anterior pole is on the left side of the frame. http://xfer.curie.fr/get/dLGUa7idKhe/mov2.avi

### Movie S3. Live imaging of transcription dynamics of *hb-MS2ΔZelda*

Expression of the *hb-MS2 AZelda* reporter in wild-type embryos. The movie has two channels: MCP-GFP channel (green) for the monitoring of nascent mRNA production dynamics and NUP-RFP (red) for nuclei detection. The capture frame is from −25% to 25% of embryo length. The anterior pole is on the left side of the frame. http://xfer.curie.fr/get/K8LogTf7AKQ/mov3.avi

### Movie S4. The transcription pattern dynamics in nuclear cycle 13

The movie shows the transcription patterns, represented by the probability of spot appearance Pspot along AP axis, at a given time after the onset of nuclear interphase. The data are from MS2 movies of *hb-MS2ΔZelda* reporter expression in wild-type embryos. Data are shown for each of the 4 individual embryos (color lines in left panel) and pulled (dashed blue line with error bars in right panel). Also shown is in the transcription pattern extracted from FISH (dashed black line with error bars) from Porcher *et al.* (2010). http://xfer.curie.fr/get/KYyBbweDQl1/mov4.avi

### Supporting Information text

http://xfer.curie.fr/get/FKDccxO73Mm/SIPDF.pdf

**Support Information**

## I. Supplementary material and methods

### BAC recombineering

the *24xMS2* cassette was first cloned into a pL-452-N-eGFP plasmid (Addgene #19173) in 3’ of a lox-Kan-lox selection cassette. Second, ~100 bp primers homologous to flanking regions of either the 5’-UTR or 3’-UTR regions of *hb* were generated to create recombination sites (respectively called 5’HR and 3’HR) and used to amplify 5’Fw-24xMS2-lox-KAN-lox-3’Rv PCR products. These PCR products were gel purified and electroporated into SW102 *E. coli* strain containing the *hb-18kb-BAC* (which contains a Chloramphenicol selection cassette) for recombination. The recombinant colonies were selected on Kanamycin (Kan) and Chloramphenicol (Cm) plates and the recombinant BAC was purified and electroporated into SW106 *E. coli* strains that carry an L-arabinose-inductible Cre gene for Kan selection cassette removal. The Kan^S^ Cm^R^ recombinant colonies were picked. The integrity of the MS2 loops sequence and the junctions of the insert at the recombination sites were verified by PCR, digestion and sequencing. The three BACs (*hb-18kb-BAC, 5’MS2-18kb-BAC* and *3’MS2-18kb-BAC*) were inserted by BestGene at the VK18 recombination site on the chromosome 2R (# 9736, Bloomington) [1].

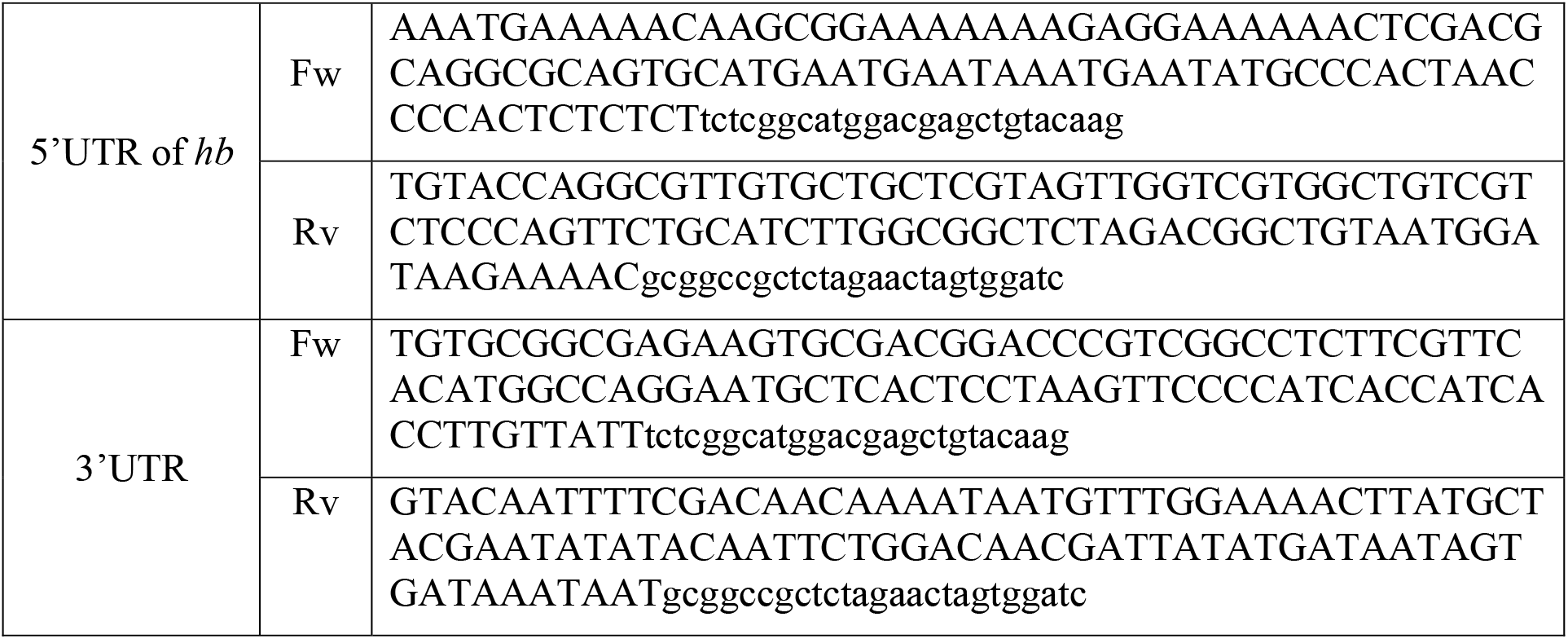
HR Primers used for recombination within the 5’UTR or 3’UTR of *hb*.

Upper cases are homologous to *hb* sequence and lower cases are homologous to pL-452-N-eGFP plasmid

### Sequence of the primers used to remove the Zelda binding site from the iRFP conding sequence

forward primer:AAAAAGATCTatggcgcgtaaggtcgatctcacctcctgcgatcgcgagccgatccac

atccccggcagcattcagccgtgcgg<u>ctgtctc</u>ctagcctgcgacg, with the mutated Zelda binding site underlined; reverse primer: AAAAAGGATCCttagcgttggtggtgggcggcggtgaagtgc).

### Sequence of the ΔZelda MS2 cassette

5’GATCCTACGGTACTTATTGCCAAGAAAGCACGAGCATCAGCCGTGCCTCAATGTCGAATCTGCAAACGACGACGATCACGCGTCGCTCCAGTATTCCAGGGTTCATCAGATCCTACGGTACTTATTGCCAAGAAAGCACGAGCATCAGCCGTGCCTCAATGTCGAATCTGCAAACGACGACGATCACGCGTCGCTCCAGTATTCCAGGGTTCATCAGATCCTACGGTACTTATTGCCAAGAAAGCACGAGCATCAGCCGTGCCTCAATGTCGAATCTGCAAACGACGACGATCACGCGTCGCTCCAGTATTCCAGGGTTCATCAGATCCTACGGTACTTATTGCCAAGAAAGCACGAGCATCAGCCGTGCCTCAATGTCGAATCTGCAAACGACGACGATCACGCGTCGCTCCAGTATTCCAGGGTTCATCAGATCCTACGGTACTTATTGCCAAGAAAGCACGAGCATCAGCCGTGCCTCAATGTCGAATCTGCAAACGACGACGATCACGCGTCGCTCCAGTATTCCAGGGTTCATCAGATCCTACGGTACTTATTGCCAAGAAAGCACGAGCATCAGCCGTGCCTCAATGTCGAATCTGCAAACGACGACGATCACGCGTCGCTCCAGTATTCCAGGGTTCATCAGATCCTACGGTACTTATTGCCAAGAAAGCACGAGCATCAGCCGTGCCTCAATGTCGAATCTGCAAACGACGACGATCACGCGTCGCTCCAGTATTCCAGGGTTCATCAGATCCTACGGTACTTATTGCCAAGAAAGCACGAGCATCAGCCGTGCCTCAATGTCGAATCTGCAAACGACGACGATCACGCGTCGCTCCAGTATTCCAGGGTTCATCAGATCCTACGGTACTTATTGCCAAGAAAGCACGAGCATCAGCCGTGCCTCAATGTCGAATCTGCAAACGACGACGATCACGCGTCGCTCCAGTATTCCAGGGTTCATCAGATCCTACGGTACTTATTGCCAAGAAAGCACGAGCATCAGCCGTGCCTCAATGTCGAATCTGCAAACGACGACGATCACGCGTCGCTCCAGTATTCCAGGGTTCATCAGATCCTACGGTACTTATTGCCAAGAAAGCACGAGCATCAGCCGTGCCTCAATGTCGAATCTGCAAACGACGACGATCACGCGTCGCTCCAGTATTCCAGGGTTCATCAGATCCTACGGTACTTATTGCCAAGAAAGCACGAGCATCAGCCGTGCCTCAATGTCGAATCTGCAAACGACGACGATCACGCGTCGCTCCAGTATTCCAGGGTTCATCA-3’

## II. Examples of MS2-MCP Time traces

**Figure S1:**
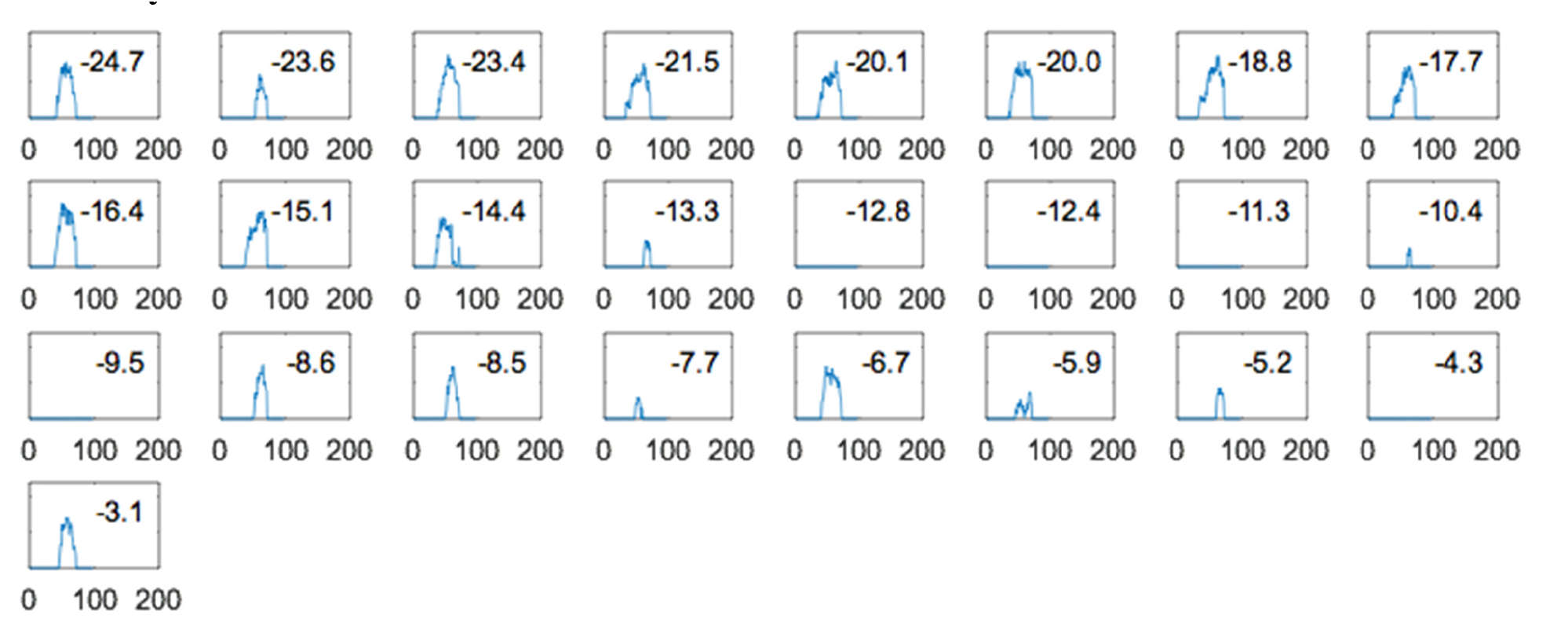
Examples of individual spot intensity over time in nuclear cycle 11. The time traces are sorted by their respective nuclei position, which is shown in the boxes in % EL (Position 0 corresponding to the middle of the embryo). Nuclei with position beyond −3.1% EL (last shown nucleus) have no spots. Horizontal axis: time in seconds. Vertical axis: spot fluorescent intensity in arbitrary units.

**Figure S2:**
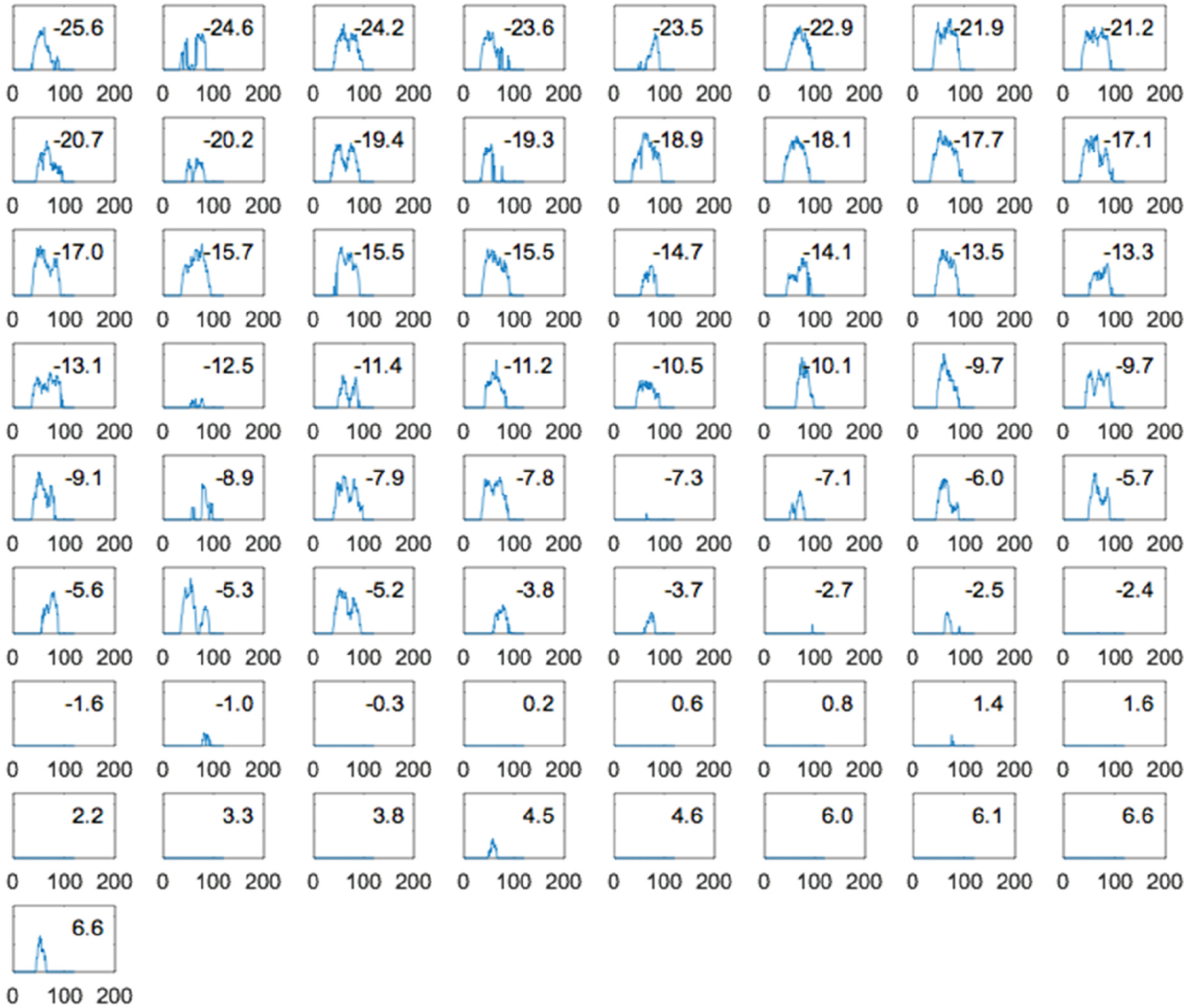
Examples of individual spot intensity over time in nuclear cycle 12. The time traces are sorted by their respective nuclei position, which is shown in the boxes in % EL (Position 0 corresponding to the middle of the embryo). Nuclei with position beyond 6.6% EL (last shown nucleus) have no spots. Horizontal axis: time in seconds. Vertical axis: spot fluorescent intensity in arbitrary units.

**Figure S3:**
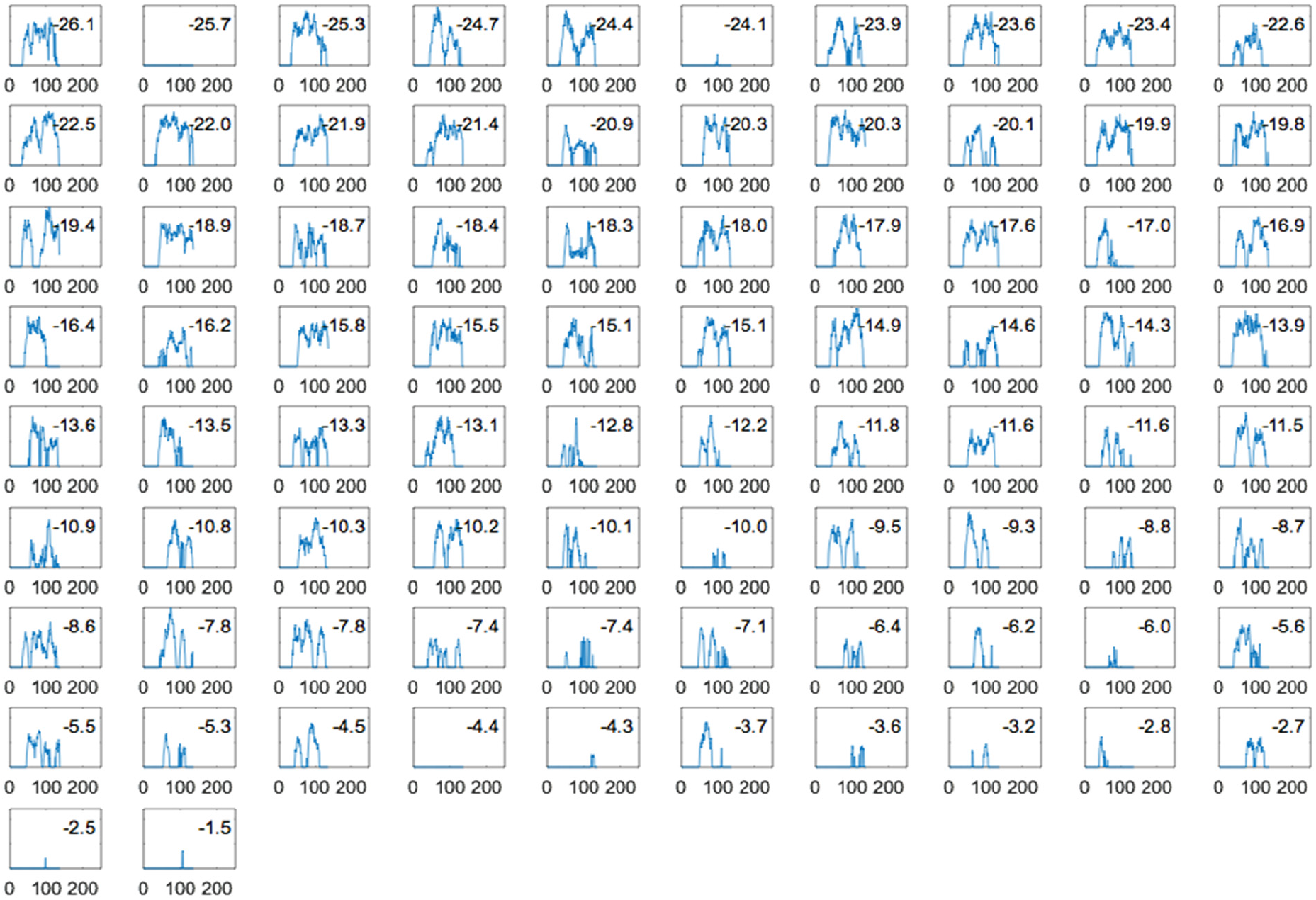
Examples of individual spot intensity over time in nuclear cycle 13. The time traces are sorted by their respective nuclei position, which is shown in the boxes in % EL (Position 0 corresponding to the middle of the embryo). Nuclei with position beyond −1.5% EL (last shown nucleus) have no spots. Horizontal axis: time in seconds. Vertical axis: spot fluorescent intensity in arbitrary units.

## III. Determining the origin of time for each time trace

During mitosis, nuclei divide in waves, starting from the two embryo poles [2]. Therefore, nuclei at the anterior pole may produce MCP-MS2 spots earlier due to either earlier chromatin decondensation or earlier reentrance of Bcd into the nucleic space. To correct for this, we first identify the birth moment of each nucleus after mitosis. A nucleus’ birth is defined as the time when the segregation from its sibling is complete (Fig. S4A-B).

**Figure S4:**
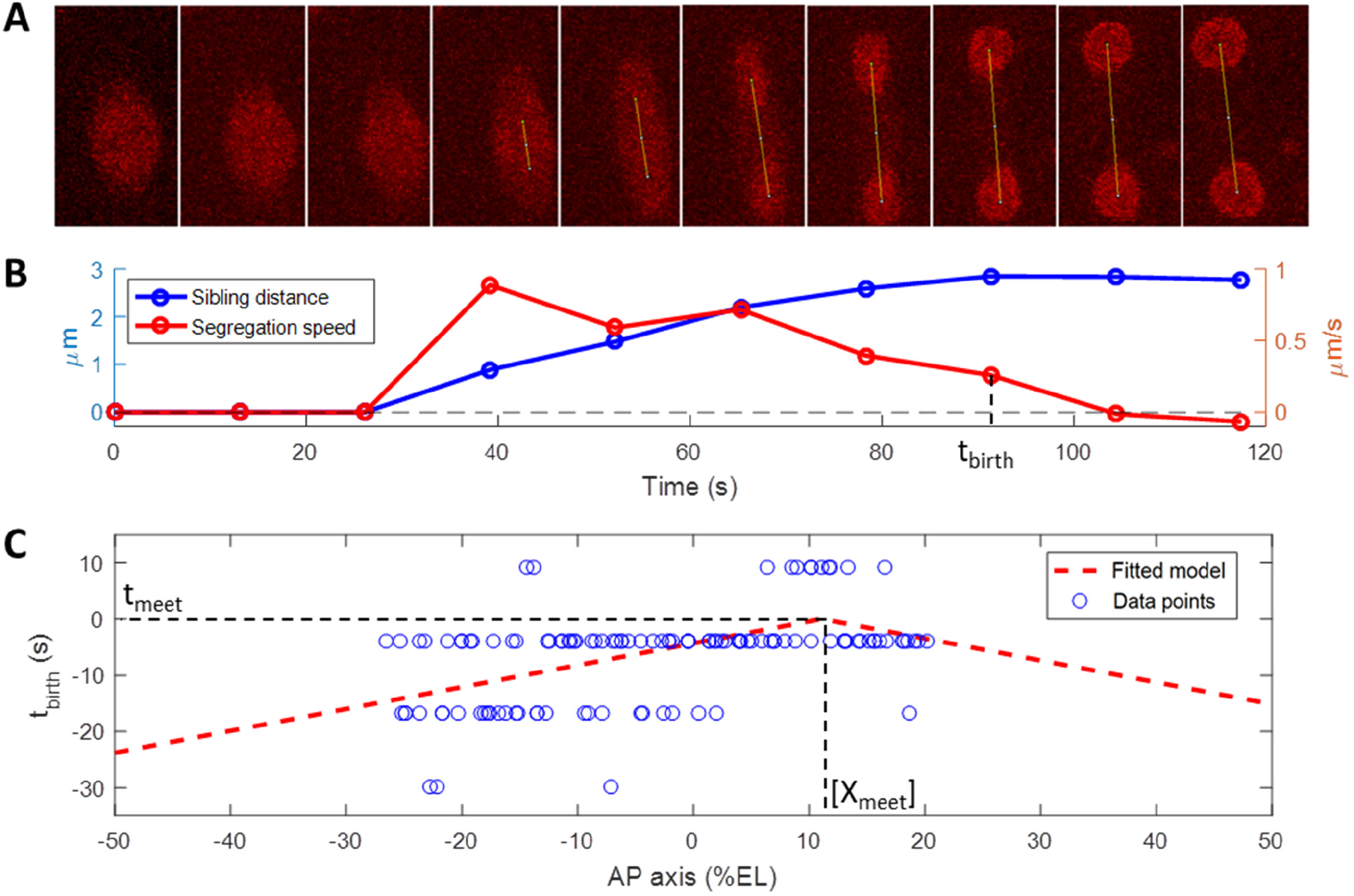
Determining the nuclei birth time: (A) Examples of frame-by-frame monitoring of two sibling nuclei after mitosis. The time interval between frames is 13.05 s. The yellow line is drawn automatically to connect the two siblings’ centroids once division is detected. (B) The distance between the centroids (blue line) and its derivative (red line) over time. The nuclei’s birth is set as the time when the speed of segregation between the sibling nuclei decreases to near-zero. (C) Examples of the nuclei birth time *t_birth_* along AP axis. Shown is the *t_birth_* extracted from the movies (blue circles) and from the fitted model in Eq. 1 (red dashed line). The y axis (*t_birth_*) is shifted so as the two mitotic waves from the two poles meet at *t_meet_*=0

To characterize the mitotic waves, we fit the birth times *t_birth_* of nuclei with siblings at position *X*along the AP axis in each embryo and nuclear cycle with a mirrored linear function:

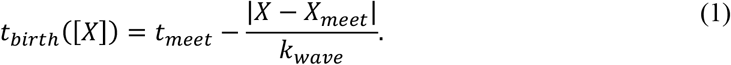

with *t_meet_* and *X_meet_* the moment and position two mitotic waves from the two poles meet. *k_wave_* (in units of % EL/s) is the speed at which the waves propagate from the poles. An example of the fit is shown in Fig. S4C. From 8 embryos, we infer *k_wave_* ~ 2.25+1.52 % EL/s, *X_meet_* ~ 8.5+15.7 % EL from mid-embryo, corresponding to the wave duration of ~ 20 s.

The intensity traces in all nuclei are then trimmed so as they start at their respective nuclei birth time. The nuclei birth times are either calculated directly for nuclei with tracked siblings or inferred through the fitted *t_birth_* for nuclei without tracked siblings (e.g. when nuclei move out of the imaging region).

## IV. Embryo alignment

Potential variations in the amount of maternal *bicoid* mRNA in the embryo’s anterior pole may lead to variations in Bicoid concentrations at a given position in different embryos and thus variations in boundary position of the expression pattern. We seek to mitigate these variations by aligning the embryos’ AP axis by their respective border position.

In order to do this, we first determine for each nucleus its “activity” feature (P_active_), which takes the value of 0 if the nucleus does not produce a MS2-MCP spot during its lifetime or 1 otherwise. We then calculate the probability for a nucleus at a given position along the AP axis to have experienced transcription of the MS2 reporter (P_ON_) as the mean of P_active_ of nuclei localized at this position The boundary position is set by inspecting where along the AP axis Pon equals 0.5. The embryos’ AP axes are then shifted to have their respective Pon border at position 0% EL.

Note that if another feature is used as the reference for the alignment (i.e. t_active_, μI or ΣI), the difference in embryos’ relative shift does not exceed 2% EL (see Table S1) when compared to P_active_.

All figures in the main manuscript use data from aligned embryos.

## V. Quantifying the steepness of *hb* expression pattern

We quantify the steepness of the *hb* expression pattern through the features of the time traces along the AP axis: the time period during which the locus is activated (t_active_), the integral transcription activity (ΣI) and the mean transcription rate (μI) (as defined in Fig. 6A of the manuscript) and Pactive (defined in SI, Section IV).

The features are first normalized by their expected value at the anterior pole to remove variations in the embryo’s growth rate and differences coming from the data acquisition process. The steepness of the feature pattern is obtained by least-square fitting the feature value along the AP axis in each embryo and nuclear cycle with a sigmoid function *f(X)*:

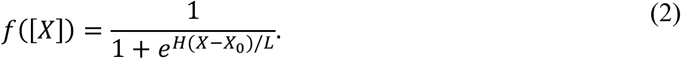

In Eq. 2, *X* is the nuclei’s position. *X_0_* and *H* are the pattern’s border position *f(X)*=0.5) and the pattern steepness (i.e. the Hill coefficient) for the feature of interest. *L* is the decay length of the Bcd gradient, which is ~100 μm or 20 % EL [3].

The results of the fit are shown in Fig. S5 (nc11), Fig. S6 (nc12), Fig. S7 (nc13) and in Table S1. In nc11, due to the very short interphase duration, the pattern is barely stabilized before mitosis, leading to very large errors in the estimated *H* (Table S1).

**Figure S5:**
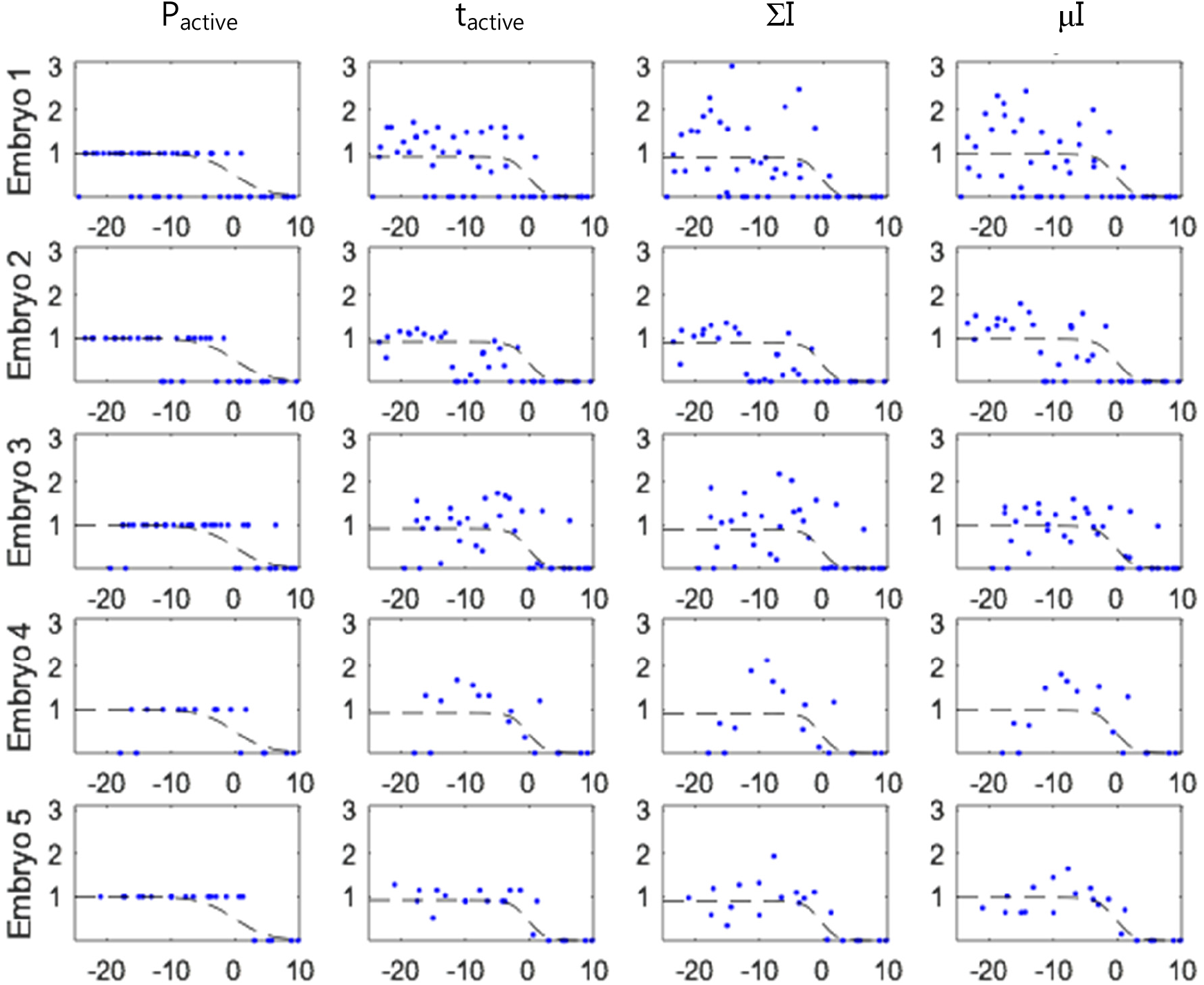
Fitting the trace feature patterns in nuclear cycle 11. The fitted curves (dashed black lines) are shown with data points (blue dots). Each data point corresponds to a single trace feature value. The horizontal axis is the AP axis in % EL.

**Figure S6:**
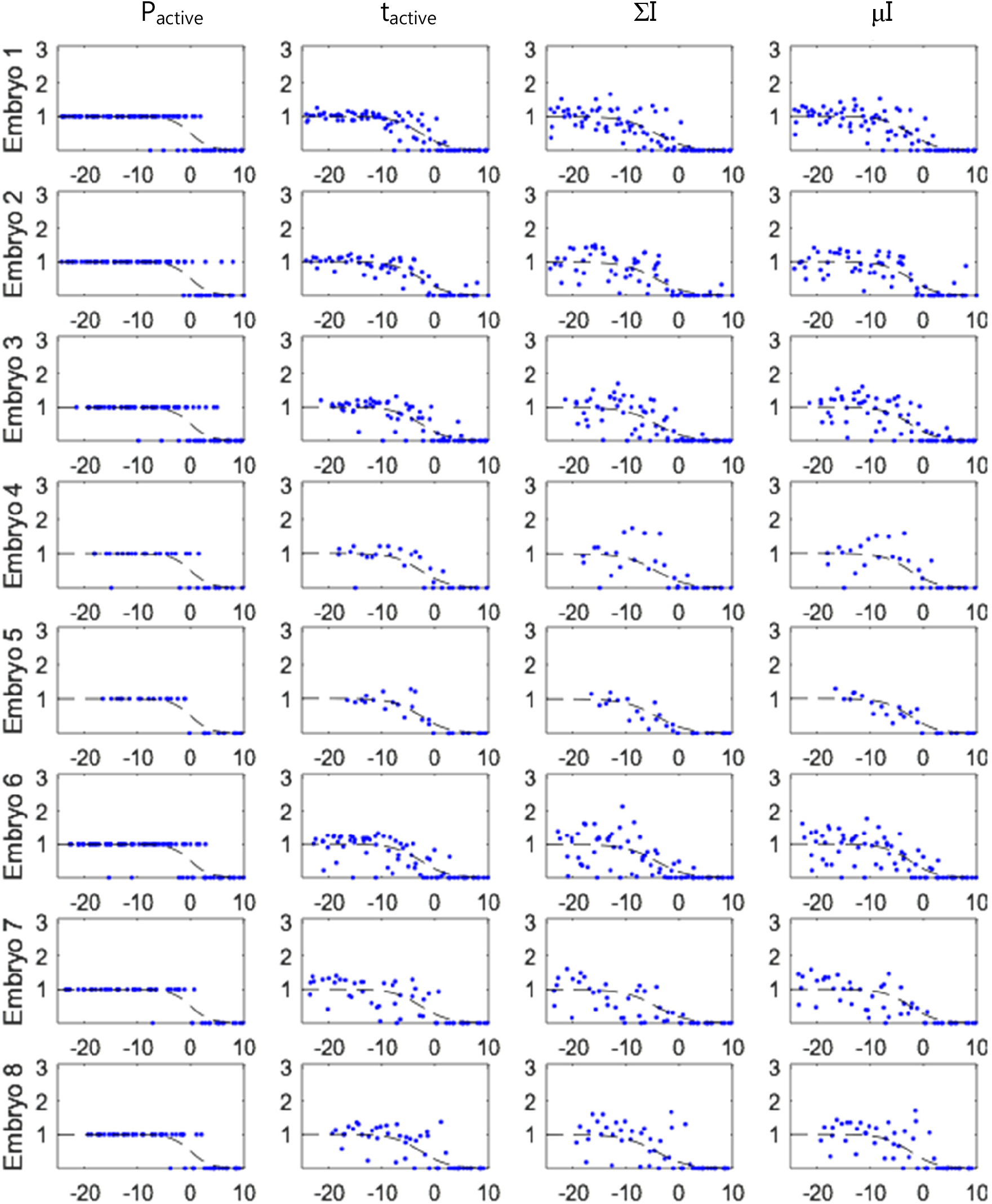
Fitting the trace feature patterns in nuclear cycle 12. The fitted curves (dashed black lines) are shown with data points (blue dots). Each data point corresponds to a single trace feature value. The horizontal axis is the AP axis in % EL.

**Figure S7:**
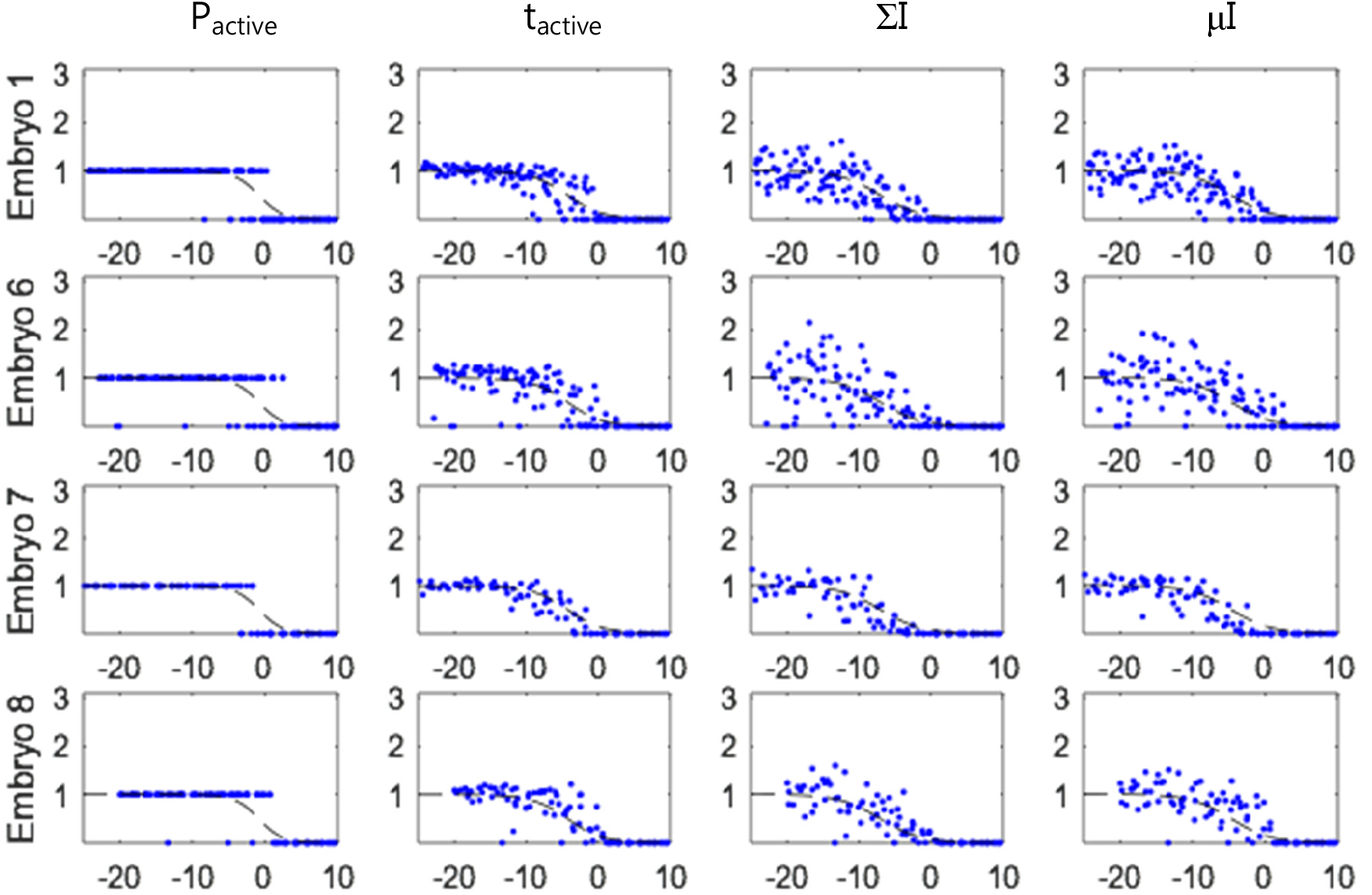
Fitting the trace feature patterns in nuclear cycle 13: The fitted curves (dashed black lines) are shown with data points (blue dots). Each data point corresponds to a single trace’s feature value. The horizontal axis is the AP axis in % EL.

**Table S1.** Parameters of fitting the sigmoid function with time trace features in nc11-13: the position of the feature pattern border (*X_0_*, in units of % EL), the pattern steepness (*H*) and their respective confidence interval (in brackets) for the locus activity (P_active_), the time period during which the locus is activated (t_active_), the integral transcription activity (ΣI) and the mean transcription rate (μI). The data are inferred from all aligned embryos in the respective cycles.

| Nuclear cycle | Feature | $P_{\text{active}}$ | $t_{\text{active}}$ | $\Sigma I$ | $\mu I$ |
| --- | --- | --- | --- | --- | --- |
| 11 | $X_0$ (% EL) | -0.1 (-1.2;1) | -0.2 (-1.2;0.8) | -0.5 (-1.8;0.8) | -0.3 (-1.4;0.8) |
| | $H$ | 7.7 (4.2;14.0) | 16.0 (8.1;∞) | 17.2 (7.2; ∞) | 15.0 (7.8; ∞) |
| 12 | $X_0$ (% EL) | 0 (-0.3;0.3) | -2.7 (-3.1;-2.3) | -4.3 (-4.9;-3.7) | -2.5 (-3.0;-2.0) |
| | $H$ | 11.2 (9.2;13.9) | 7.8 (6.6;9.3) | 6.9 (5.8;8.5) | 8.1 (6.7;10.1) |
| 13 | $X_0$ (% EL) | -0.9 (-1.2;-0.6) | -4.3 (-4.6;4.0) | -7.0 (-7.5;-6.5) | -4.7 (-5.2;-4.3) |
| | $H$ | 12.6 (10.7 15.0) | 8.2 (7.2;9.3) | 7.1 (6.2;8.3) | 7.3 (6.3;8.4) |

### VI. Evolution of the transcription pattern steepness over time

At a given time, the *hb* pattern is characterized by the snapshot proximal *hb* promoter’s activity P_bright_, which takes value of 0 if the nucleus has no active locus and 1 if the nucleus has an active locus at the given time. At a given position, P_bright_ has a mean value of P_SPOT_ (as described in the Main manuscript). We fit the values of P_bright_ along the AP axis with a sigmoid function and infer on the pattern steepness H(t) as a function of time (Fig. S8) for each nuclear cycle.

**Figure S8.**
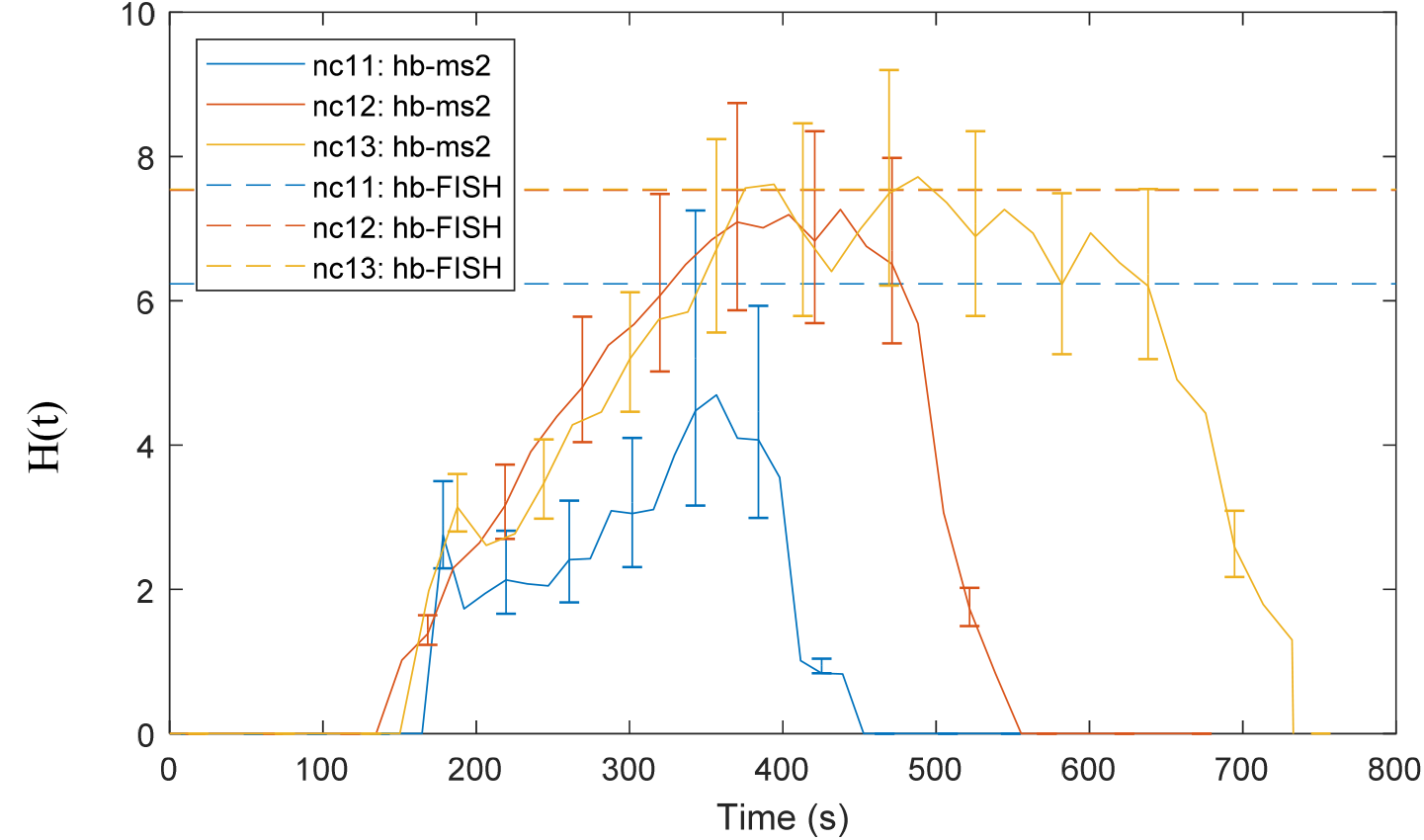
Time evolution of the pattern steepness H(t) over time: shown for nc 11 (blue solid line), nc 12 (red solid line) and nc 13 (yellow solid line) along with the margins of errors (p-value = 0.05). Also shown (dashed lines) are the Hill coefficients extracted from FISH data in for the respective cycles. The coefficients from FISH in nc12 and nc13 are almost identical.

### VII. Models of transcription factor (TF) sensing and transcription

#### Model of TF sensing

We assume the previously suggested [4–6] model of transcription factor sensing through the multiple TF binding sites:

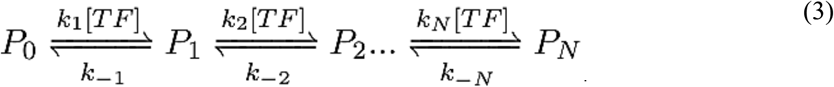

*N* is the number of operator sites for TF binding. *P_i_* is the promoter state when bound by *i* transcription factor (TF) molecules. The operator sites are assumed to be identical, thus do not need to keep track of which operator sites are currently occupied in *P_i_. [TF]* is the TF concentration normalized by the TF concentration at the boundary position *X_0_*. In the case of the Bcd-hb system, *[TF]* follows a decaying gradient from the anterior with the decay length L ~ 100 μm or 20 % EL [3]:

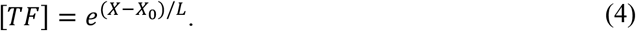

*k_i_* and *k_−i_* are the binding and unbinding rate constants respectively. We assume that there is no cooperativity in the searching of the TF for the operator sites: when the promoter is at state *P_i-l_*, TFs can bind to any of the remaining free operator sites independently at a rate:

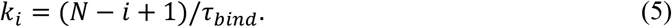

*τ_bind_* is the time for a free operator to be bound by any TF at the *hb* expression boundary (*X=X_0_, [TF]*=1). Assuming that the TF can only search for OS by diffusing in the nucleus and that each collision between TF and OS is one successful binding event, *τ_bind_* depends on the diffusion coefficient of the TF (*D* ~ 7.7 μm^2^/s [7] the absolute TF concentration at the boundary (*c* ~ 11.2 molecules/μm^3^) [8] and the size of the operator site (*a* ~ 0.003 μm [6]).

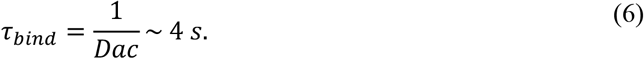

We call *P(P_i_,X, t)* the probability of the promoter to be in state *P_i_* at time t and position *X*. The time evolution of *P(P_i_,X,t)* is given by:

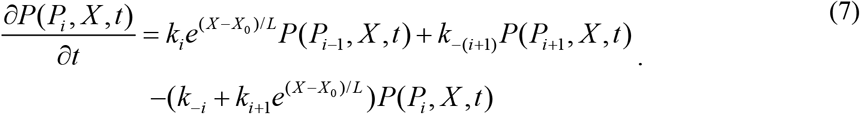

For convenience, we define the effective equilibrium constants 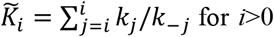 and 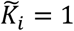 for i=0. By solving the linear differential equations corresponding in Eq. 7, we have the probability of the promoter in state *P_i_* at steady state at a given position *X* [9]:

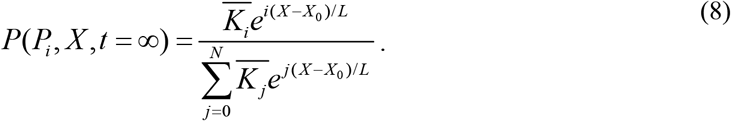

In this work, we consider the “all-or-nothing” case: the *hb* gene becomes active only when all the OS of the *hb* promoter are bound (*P_active_ ≡ PN*).

#### 2) Transcription initiation

While the gene is activated (i.e. the promoter is fully bound by TF), RNAP can bind stochastically to the promoter and initiate transcription at a rate λ ~ 0.15 s^−1^. In the initiation process, it occupies the promoter for a duration *t_biock_ ~ 6 s* [10] preventing the binding of another RNAP to the promoter until the RNAP frees the promoter site. The values of *λ* and *τ_biock_* are extracted from the transcription dynamics of *hb* at the anterior pole [11].

#### 3) Transcription elongation

Promoter escape is followed by a deterministic transcription elongation process [12] with a rate constant 40bp/s [13]. During this process, MCP-GFP can quickly bind to the newly transcribed MS2 binding sites on the nascent RNA.

We denote *L(t)* the number of MS2-MCP binding sites on a nascent RNA at time t after its transcription initiation. *L(t)* depends only on the reporter gene construct. In this work, with the ms2 binding site array located at the 3” end of the reporter gene, *L(t)* is given in Fig. S9.

**Figure S9:**
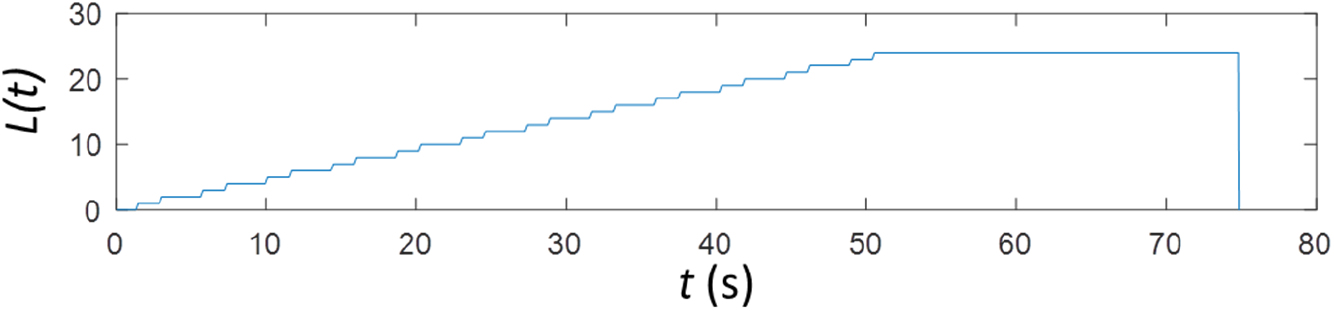
MS2 binding site configuration. *L(t) is* the number of MS2-MCP binding sites on a nascent RNA at time *t* after its transcription initiation.

The intensity of the transcription loci is therefore given by the convolution of the transcription initiation signal *I_RNAP(t)_* with the reporter gene configuration function *L(t)*:

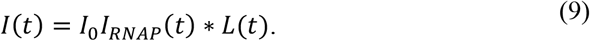

Here, *I_0_* is the intensity of a single bound MS2-MCP binding site.

#### 4. The pattern steepness and the promoter dynamics out of steady state

For a given set of kinetic parameters 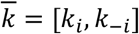 in Eq. 3, we quantify the pattern steepness and the evolution of the promoter mean activity over time. We only focus on the boundary position (*X=X_0_*) where the gene is 50% activated at steady-state (*P(P_active_, X_0_*,∞)=0.5).

##### Pattern steepness

We calculate the pattern steepness *H* from the slope of the promoter activity pattern at the boundary position (*X=X_0_*) at steady state:

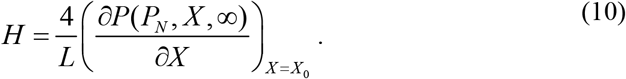

##### Promoter dynamics of steady state

The vector *s(t)=[P(P_0_,X,t), P(P_0_,X,t)*,… *P(P_0_,X,t)*]^T^ describes the probability for nuclei at a position X and time to be in a given promoter state. The change in the vector *s(t)* is described by the transition matrix *U*, whose elements are defined by the stochastic equations in Eq. 7 evaluated at the boundary position (*X=X_0_*). At time *t=0*, we assume that the promoter is free of TF: s(0)=[1,0…,0]^T^ and the TF interact with OS of the promoter. The mean promoter activity at time *t* is given by:

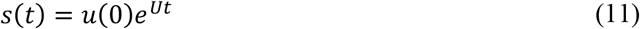

We define a vector α indicating which promoter state is associated with the target being activated. In the case of “all-or-nothing” (*P_active_ =* P_N_), α=[0,0…,1]^T^. The distribution of nuclei with active loci is given by:

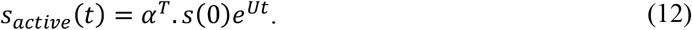

##### Fitting the transcription pattern

We find the kinetic parameter set 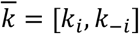 that matches the observed Hill coefficient and the promoter state probability at the boundary (*X=X_0_*) at near steady state (330 s from the beginning of nuclear cycle or ~180 s after first spot appearances). For simplicity, we approximate the probability of the activated gene *s_active_*(t=180 s) by the probability of the spot appearance in each nuclei P_SPOT_ at 330 s following mitosis.

From the measurements, we infer the Hill coefficient H ~ 7.1 + 0.53, *s_active_*(180 s) ~ 0.45 + 0.02.

We fit the model’s Hill coefficient and the promoter active state distribution to the data using least square fit, weighted by the margin of error. The value of 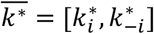 can be found by minimizing the objective:

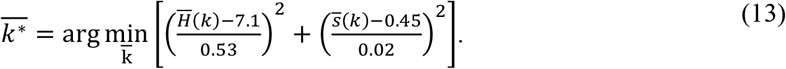

Where 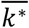 and 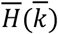 are respectively the steepness and the probability of activated gene calculated numerically from Eq. 10 and 12 for any given parameter set 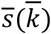

Due to the model complexity and the high number of parameters involved with 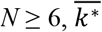 is obtained by a brute force search, followed by local optimization (~ 10^5^ iterations for each value of *N*). The unbinding rates *k_−i_* are randomized from 10^−20^ s^−1^ to 10^20^ s^−1^ while the binding rates *k_i_* are kept constant (Eq. 5).

The fitted kinetic parameters are shown in Table S2.

**Table S2:** Fitted models of varying number of operator sites *N* and the model of 6 independent sites. Shown is the p-value of the likelihood ratio test between the fitted model of *N* and *N-1* OS. Also shown is the Bayesian Information Criterion (BIC) for each model. The data for the fitting is pulled from all embryos in all nuclear cycles.

| $\bar{k}^*$ | $N=6$ | $N=7$ | $N=8$ | $N=9$ | $N=10$ | $N=6$<br>*Ind.* |
| --- | --- | --- | --- | --- | --- | --- |
| $k_{-1} \text{ (s}^{-1}\text{)}$ | 21.34 | 31.02 | 1.12 | 4.20 | 0.03 | 0.48 |
| $k_{-2} \text{ (s}^{-1}\text{)}$ | 1.50 | 0.92 | 16.15 | 8.14 | 36.46 | 0.97 |
| $k_{-3} \text{ (s}^{-1}\text{)}$ | 0.51 | 0.51 | 1.87 | 2.01 | 0.39 | 1.46 |
| $k_{-4}$ (s<sup>-1</sup>) | 2.47 | 1.60 | 2.12 | 1.59 | 4.33 | 1.94 |
| $k_{-5}$ (s<sup>-1</sup>) | 0.19 | 0.82 | 1.25 | 1.20 | 1.59 | 2.43 |
| $k_{-6}$ (s<sup>-1</sup>) | 0.016 | 0.84 | 0.31 | 0.47 | 3.10 | 2.91 |
| $k_{-7}$ (s<sup>-1</sup>) | / | 0.011 | 0.80 | 1.85 | 0.40 | / |
| $k_{-8}$ (s<sup>-1</sup>) | / | / | 0.013 | 0.19 | 0.76 | / |
| $k_{-9}$ (s<sup>-1</sup>) | / | / | / | 0.025 | 0.31 | / |
| $k_{-10}$ (s<sup>-1</sup>) | / | / | / | / | 0.023 | / |
| $H(\bar{k})$ | 5.15 | 5.48 | 6.11 | 6.56 | 6.69 | 1.31 |
| $s(\bar{k})$ | 0.43 | 0.45 | 0.43 | 0.44 | 0.44 | 0.48 |
| p-value | 1 | 0.0050 | 0.0044 | 0.0133 | 0.645 | / |
| BIC | 45.7 | 40.7 | 35.4 | 32.1 | 24.7 | 212.6 |

Given the p-value of the likelihood ratio test between the models and the Bayesian Information Criterion (BIC) for each model, we find that increasing *N* up to 9 results in significant better fit of the model with data. Increasing *N* beyond 9 does not improve the fit significantly.

##### 6) Stochastic simulations

We use the Stochastic Simulation Algorithm [14, 15] to simulate the promoter dynamics under the regulation of the TF and the timing of the transcription initiation events by RNAP *I_RNAP_(t)*. The trajectory of *I_RNAP_(t)* is then convoluted with the gene configuration function *L(t)* to achieve the spot intensity *I(t)* over time. An example of the intensity trace is shown in Fig. S10.

**Figure 10.**
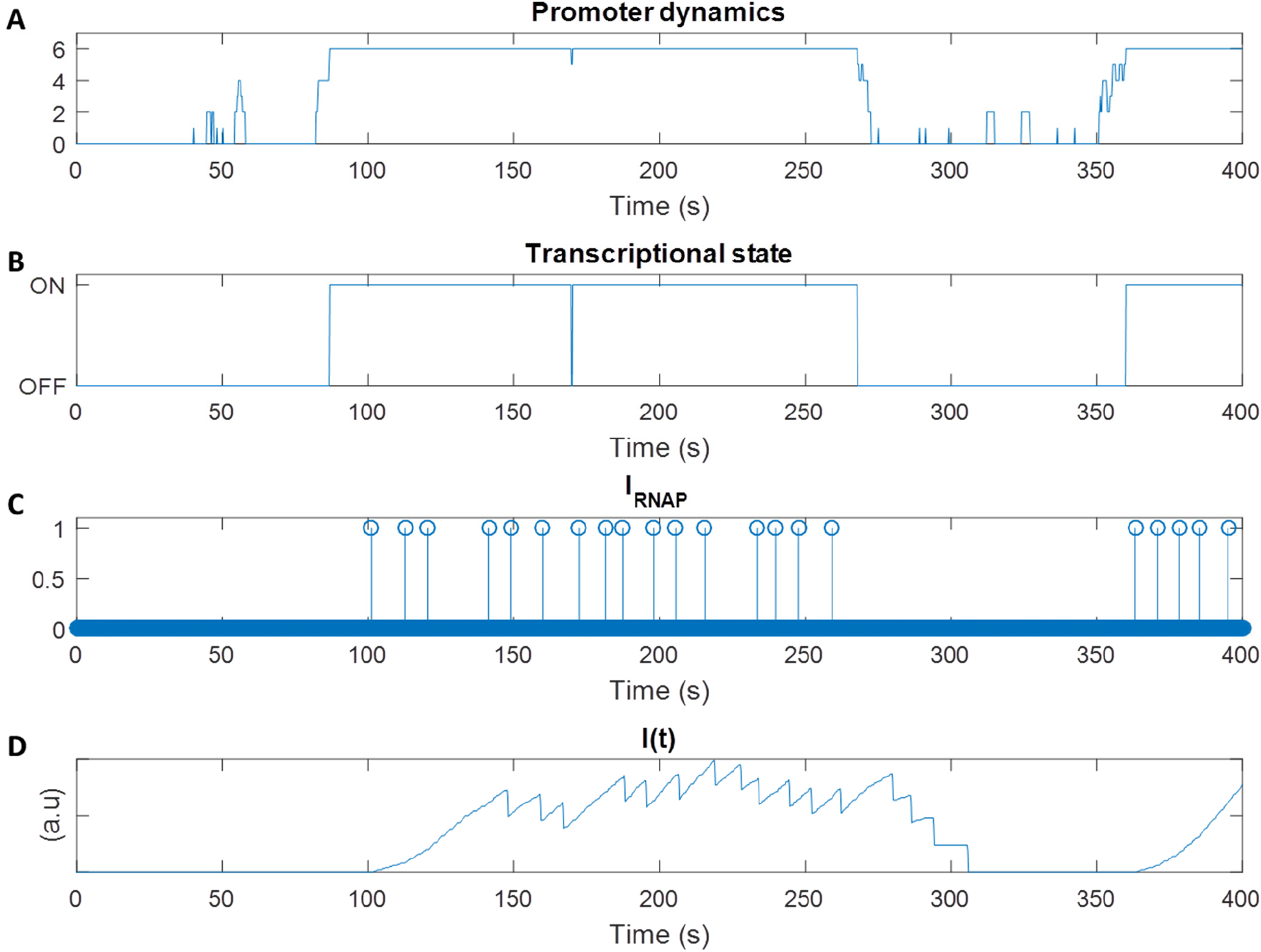
Examples of simulated trajectories at mid-embryo for *N=6.* (A) The number of bound TF molecules to the promoter over time. (B) The gene transcriptional state given the number of bound molecules to the promoter. The gene is turned ON when the promoter is fully bound by TF. (C) Occurrences of transcription initiation events *I_RNAP_(t)*, corresponding to (B). (D) Transcription loci intensity *I(t)*, corresponding to (C).

At each position along the AP axis, with the fitted kinetic parameters, we simulate 500 nuclei intensity traces, from which the heatmap of P_SPOT_ over time and along AP axis can be constructed (Fig. 8 in main Manuscript).

### VII. Comparing the noise in positional readout between models

With the fitted parameters (Table S2), we compare the precision of gene expression readout *f_T_* between the case *N*=6, *N*=9 at the boundary position (*X=X_0_*). Here, the readout is defined as the mean duration the *hb* gene is activated at steady-state:

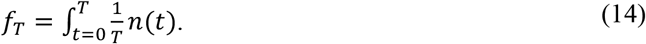

with *n(t)* being the trajectory of the gene activity state over time. *n(t)* is 1 when the gene is activated and 0 otherwise. The relative noise in the readout *CV_p_* is defined as follow:

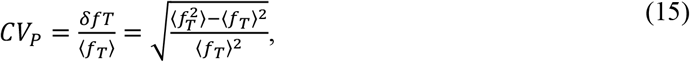

in which *<fT>*=0.5 and *<fτ^2^>* are respectively the first and second moments of the readout at the pattern’s boundary (*X=X0*). Let us define a vector s_fire_ where s_fire,i_=α_i.si._ *<fτ^2^>* is calculated numerically from the transition matrix U [9]:

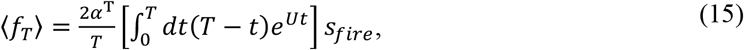

The precision of gene expression readout between the model with *N=6* and *N*=9 are shown in Fig. S11. Also shown is precision from the “no cooperativity” case, where interactions of TF with the binding sites are independent (k_−i_ = i.k_−N_/N)

**Figure S11.**
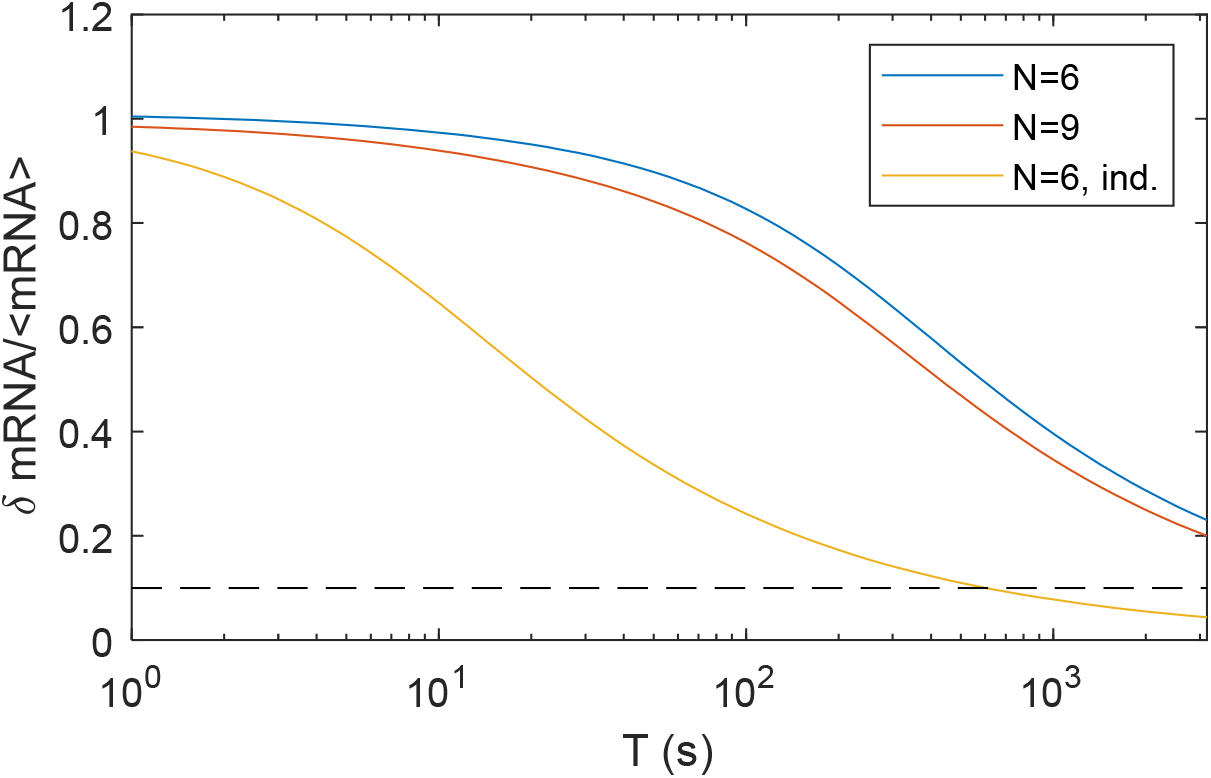
Comparing the relative noise in the positional readout *δmRNA/<mRNA>* as a function time: for fitted model with *N*=6 (blue line), fitted model with *N*=9 (red line) and “no cooperativity” model (yellow line).

